# EC1 body controls sperm cell reception *via* gamete pit formation

**DOI:** 10.64898/2026.07.14.738444

**Authors:** Daichi Susaki, Hinako Oshirabe, Shiori Nagahara, Yumi Goto, Takao Oi, Hidenori Takeuchi, Mayuko Sato, Kiminori Toyooka, Hiromi Kakizaki, Naoya Sugi, Ari Yoshimura, Kaoru Tonosaki, Tetsu Kinoshita, Daisuke Maruyama

## Abstract

The remarkable efficiency of double fertilization in flowering plants depends on the precise delivery of immotile sperm cells to two female gametes^1^. Although sperm cells are rapidly released from the pollen tube, how they are accurately directed to their fertilization site has remained unknown. Here we identify a transient extracellular structure, termed the gamete pit, that forms at the egg cell–central cell (EC–CC) interface immediately after pollen tube discharge and serves as the landing site for sperm cells. Gamete pit formation is enabled by localized dissociation of the EC–CC interface, mediated by extracellular protein assemblies designated EC1 bodies. These structures contain extracellular deposits enriched in EGG CELL 1 (EC1) peptides^2^, including amyloid-like assemblies. Loss of EC1 bodies prevented gamete pit formation, causing sperm-cell backflow or catastrophic over-penetration of pollen tube contents into the central cell. Surprisingly, ectopic deposition of a heterologous amyloidogenic protein alleviated over-penetration and partially restored fertility. Our findings reveal a mechanism by which extracellular protein assemblies remodel the gamete interface for sperm-cell reception, uncover an unexpected structural role for the conserved gamete-activating peptide EC1, and establish protein-mediated cell-interface remodeling as a fundamental principle underlying double fertilization in flowering plants.

---

In contrast to flagellate sperms in moss and fern, flowering plants (angiosperms) accomplish fertilization using non-motile sperm cells delivered by pollen tube^3^. The evolution of this male genome delivery system is thought to have enabled successful reproduction in dry terrestrial environments by eliminating the need for free water during fertilization. Pistils contain ovules, the precursors of seeds, in which two female gametes—the egg cell and central cell—are enclosed together with two synergid cells that attract the pollen tube. Upon perceiving guidance cues from the synergids, the pollen tube grows into the ovule, releases two sperm cells, and double fertilization ensues (Fig. 1a)^4^.

**Fig. 1.**
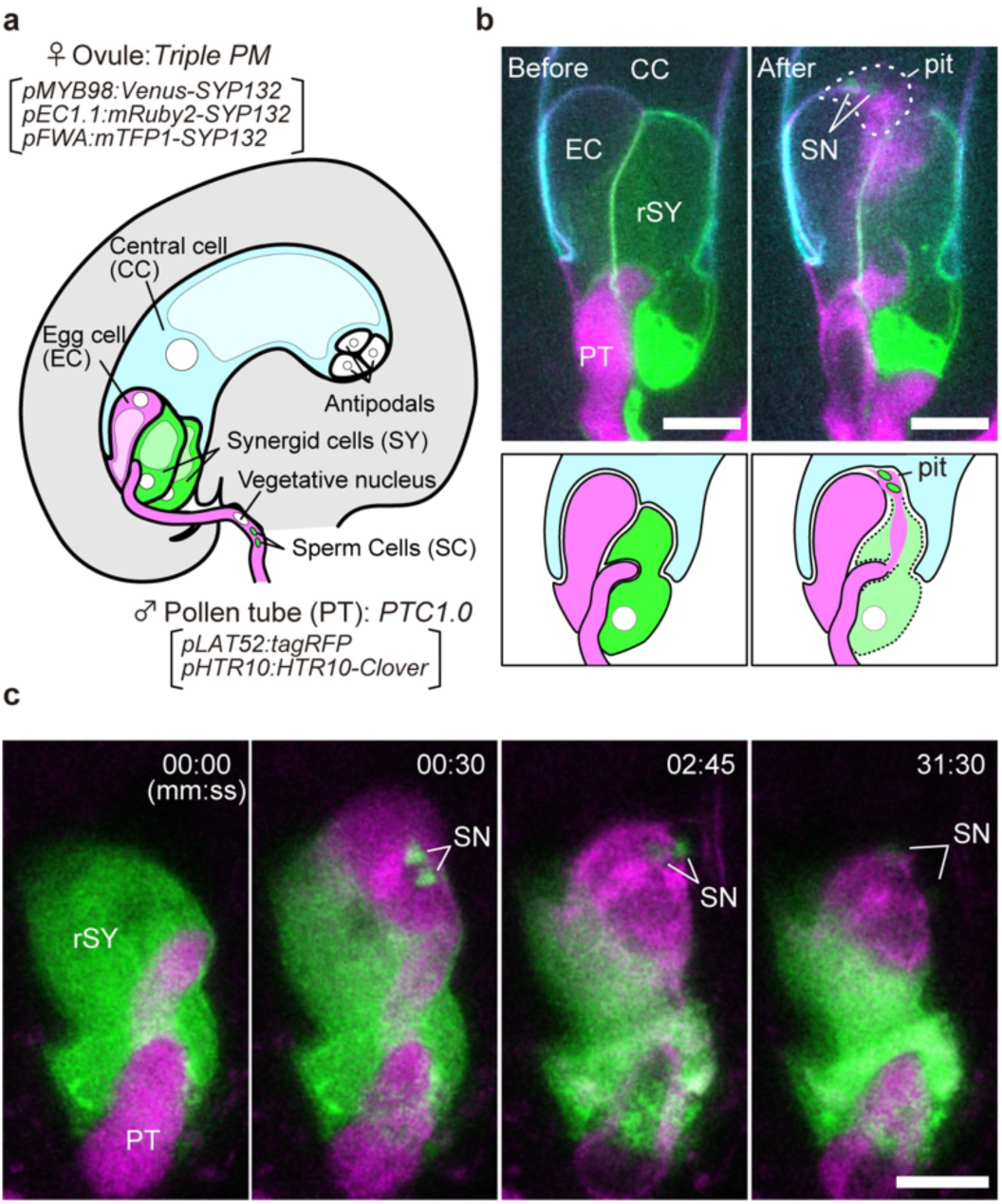
Synergid penetration and gamete pit formation upon pollen tube discharge. **a**, Schematic illustration of an *Arabidopsis* ovule immediately before pollen tube discharge. The *Triple PM* marker labels the plasma membranes of the egg cell (mRuby2), central cell (mTFP1), and synergid cells (Venus). Pollen tube contents were visualized using the *PTC1.0* marker, consisting of a pollen tube cytosolic marker (*pLAT52:tagRFP*) and a sperm nuclear marker (*pHTR10:HTR10-Clover*). **b,** Representative maximum-intensity projections from time-lapse confocal imaging before and after pollen tube discharge (top) together with corresponding schematic illustrations (bottom). Semi-*in vivo* fertilization assays were performed using ovules expressing the *Triple PM* marker and pollen expressing the *PTC1.0* marker. See also Supplementary Video 1. **c,** Time-lapse imaging of pollen tube penetration into the receptive synergid using the plasma membrane marker *pMYB98:PIP2A-GFP*. Three-dimensional reconstruction confirmed that the receptive synergid plasma membrane completely envelops the penetrating pollen tube contents throughout synergid invasion. See also Supplementary Videos 2 and 3. Abbreviations: PT, pollen tube; EC, egg cell; CC, central cell; SY, synergid cell; rSY, receptive synergid; SC, sperm cell; SN, sperm nucleus. Scale bars, 10 μm.

Although these events occur deep within the carpel and long remained inaccessible to direct observation, live-cell imaging in *Torenia fournieri* and *Arabidopsis thaliana* has revealed key steps of the fertilization process^1,5–11^. Following pollen tube discharge, sperm cells are rapidly carried by the cytoplasmic efflux and reach the EC–CC interface within seconds^1^. Remarkably, their trajectories are nearly linear, suggesting that sperm-cell release is spatially controlled rather than mechanically stochastic. However, the cellular mechanism that determines the final destination of released sperm cells has remained unknown.

Here we identify a transient extracellular structure, the gamete pit, that forms through localized dissociation of the EC–CC interface and functions as the landing site for sperm cells. We further show that this process is mediated by extracellular protein assemblies enriched in the conserved gamete-activating peptide EC1, uncovering an unexpected structural function for EC1 and revealing protein-mediated cell-interface remodeling as the mechanism underlying precise sperm-cell reception.

## Gamete pit formation

To determine how released sperm cells reach their fertilization site, we simultaneously visualized pollen tube discharge together with the plasma membranes of all female gametophytic cells during *Arabidopsis* fertilization. Based on previous live-imaging studies, Sprunck and colleagues proposed that the EC–CC interface is transiently pushed open by the force of pollen tube discharge, allowing immotile sperm cells to enter the EC–CC boundary^12^. However, because all female gametophytic cells had not previously been visualized simultaneously during pollen tube reception, the dynamics and structural basis of this proposed opening remained unresolved. We therefore combined a pollen tube content marker (*PTC1.0*), consisting of a tagRFP-labelled pollen tube cytosolic marker (*pLAT52:tagRFP*) and a Clover-labelled sperm nuclear marker (*pHTR10:HTR10-Clover*), with a *Triple PM* marker that labels the plasma membranes of the egg cell, central cell and synergid cells with mRuby2, mTFP1 and Venus, respectively (Fig. 1a). Semi-*in vivo* fertilization assays using these marker lines allowed simultaneous visualization of pollen tube discharge and membrane dynamics throughout sperm-cell reception (Fig. 1a,b and Supplementary Video 1).

Following invasive pollen tube growth, released pollen tube contents penetrated through the receptive synergid, consistent with a recent preprint^11^. Three-dimensional reconstruction using the *pMYB98:PIP2A-GFP* plasma membrane marker confirmed that the receptive synergid plasma membrane remained wrapped around the penetrating cytoplasmic stream (Fig. 1c and Supplementary Videos 2 and 3). These observations indicate that the rigid cell wall of the receptive synergid acts as a cylindrical mechanical guide for the rapidly released pollen tube contents.

Unexpectedly, the released pollen tube contents emerged beyond the receptive synergid and displaced the central-cell plasma membrane while only minimally deforming the egg-cell plasma membrane, thereby generating a distinct extracellular cavity at the EC–CC interface (Fig. 1b and Supplementary Videos 1–3). We designate this transient structure the gamete pit. Although previous studies proposed transient opening of the EC–CC interface during pollen tube discharge^11–13^, direct visualization of the membrane dynamics reveals that sperm-cell reception occurs through formation of a discrete extracellular cavity rather than along a pre-existing narrow interface. The emergence of the gamete pit suggested that the EC–CC interface undergoes a specialized structural rearrangement during pollen tube discharge, prompting us to investigate the structural basis of gamete pit formation.

## Patchy extracellular structure

The transient opening of the EC–CC interface suggested the presence of a specialized extracellular architecture that permits rapid yet localized separation during pollen tube discharge. Ultrastructural studies in diverse angiosperms, including crops and the basal angiosperm *Amborella trichopoda*, have consistently described discontinuous extracellular material along the EC–CC interface^12–18^, suggesting that this distinctive architecture is evolutionarily conserved. In mature *Arabidopsis* ovules, transmission electron microscopy (TEM) similarly revealed discontinuous extracellular patches at the EC–CC interface (Fig. 2a, arrowheads). However, because these observations were limited to individual sections, the three-dimensional organization of the extracellular patches remained unknown. We therefore reconstructed the EC–CC interface from serial focused ion beam scanning electron microscopy (FIB-SEM) images. Three-dimensional reconstruction revealed that the discontinuous profiles constitute an interconnected extracellular network extending specifically along the EC–CC interface (Fig. 2b and Supplementary Video 4). Notably, comparable structures were absent from the egg cell–synergid interface, indicating that this extracellular architecture is unique to the two female gametes.

**Fig. 2.**
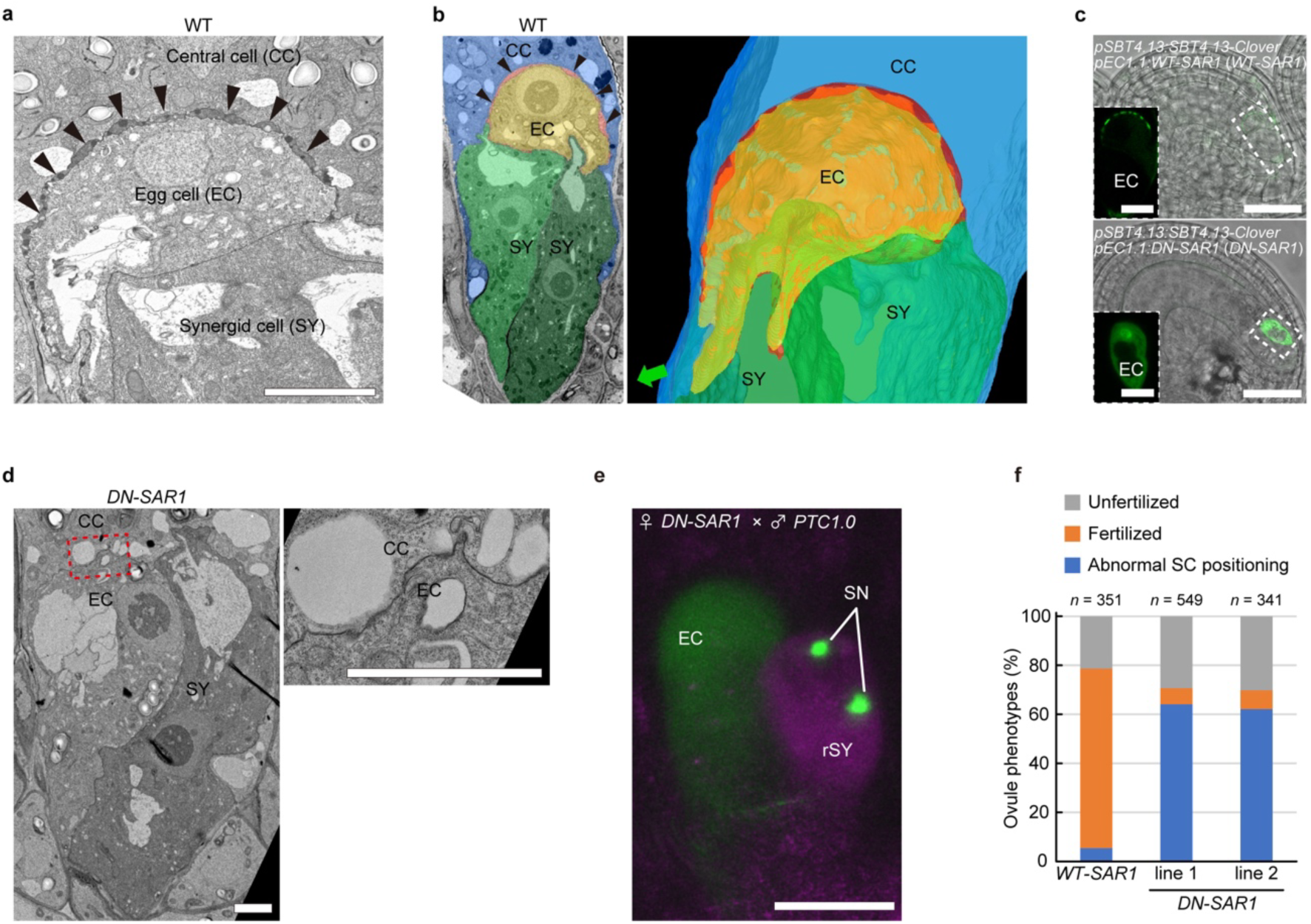
Egg-cell secretion establishes extracellular architecture at the EC–CC interface. **a**, Representative transmission electron micrograph of the EC–CC interface in a mature wild-type ovule. Arrowheads indicate discontinuous extracellular patches. Representative image from n = 9 ovules. **b,** FIB-SEM analysis of the EC–CC interface. Left, representative single section. Right, three-dimensional reconstruction of manually segmented egg cell, central cell, synergid cells, and extracellular patches generated from serial FIB-SEM sections. See also Supplementary Video 4. **c,** Localization of the egg-cell-secreted subtilase reporter *pSBT4.13:SBT4.13-Clover* in ovules expressing wild-type SAR1 (WT-SAR1) or dominant-negative SAR1 (DN-SAR1) specifically in the egg cell. Insets show higher-magnification views of the boxed regions. See also Extended Data Fig. 1 and Supplementary Video 5. **d,** Representative transmission electron micrographs of the EC–CC interface in *DN-SAR1* ovules (n = 4). Right, magnified view of the dashed box shown in the left panel. The EC–CC interface in DN-SAR1 ovules lacks extracellular patches and exhibits extensive close apposition of the egg-cell and central-cell plasma membranes. **e,** Representative confocal images showing sperm-cell behavior following pollen tube discharge in DN-SAR1 ovules. Released sperm cells frequently remain within the receptive synergid and localized EC–CC dissociation fails to occur. See also Supplementary Video 6. **f,** Quantification of ovules exhibiting defective sperm-cell reception in *WT-SAR1* and *DN-SAR1* plants. Ovules were analyzed 14–16 h after pollination using PTC1.0 pollen. Abbreviations: EC, egg cell; CC, central cell; SY, synergid cell; rSY, receptive synergid; SN, sperm nucleus. Scale bars, 5 µm in (**a**); 3 µm in (**d**); 10 µm in (**c**) and (**e**).

To investigate the origin of these extracellular patches, we examined *pSBT4.13:SBT4.13-Clover*, a translational fusion reporter for the egg-cell-secreted subtilase SBT4.13^19,20^. Fluorescent signals precisely marked the extracellular patches (Extended Data Fig. 1 and Supplementary Video 5), suggesting that protein secretion contributes to their formation or maintenance. Consistent with this possibility, inhibition of COPII-dependent secretion specifically in the egg cell by dominant-negative SAR1^21^ abolished SBT4.13-labelled extracellular patches, whereas expression of wild-type SAR1 had no detectable effect (Fig. 2c). TEM further revealed extensive close apposition of the egg-cell and central-cell plasma membranes in *DN-SAR1* ovules (Fig. 2d), whereas extracellular patches remained unaffected when dominant-negative SAR1 was expressed from central cell-specific promoters (Extended Data Fig. 2) ^22,23^.

To determine whether these extracellular patches are required for gamete pit formation, we analysed pollen tube reception in DN-SAR1 ovules. Following pollen tube discharge, released sperm cells frequently remained within the receptive synergid and the characteristic cavity observed in wild-type ovules failed to develop (Fig. 2e,f and Supplementary Video 6). These observations demonstrate that egg-cell secretion is required to establish the extracellular architecture that enables localized EC–CC dissociation during gamete pit formation. The molecular identity of these extracellular patches therefore became the next key question.

## EC1 body

To determine the molecular identity of the extracellular patches, we revisited the localization of the EGG CELL 1 (EC1) family, whose members have been proposed to accumulate within the egg cell before being secreted upon pollen tube discharge to activate sperm cells^2^. Translational reporters for all five *Arabidopsis* EC1 family members consistently labelled the extracellular patches before pollen tube arrival (Fig. 3a), indicating that EC1 proteins are secreted before pollen tube discharge rather than being retained exclusively within the egg cell. Representative three-dimensional reconstruction of EC1.4-Clover further revealed a continuous extracellular network extending along the EC–CC interface (Supplementary Video 7).

**Fig. 3.**
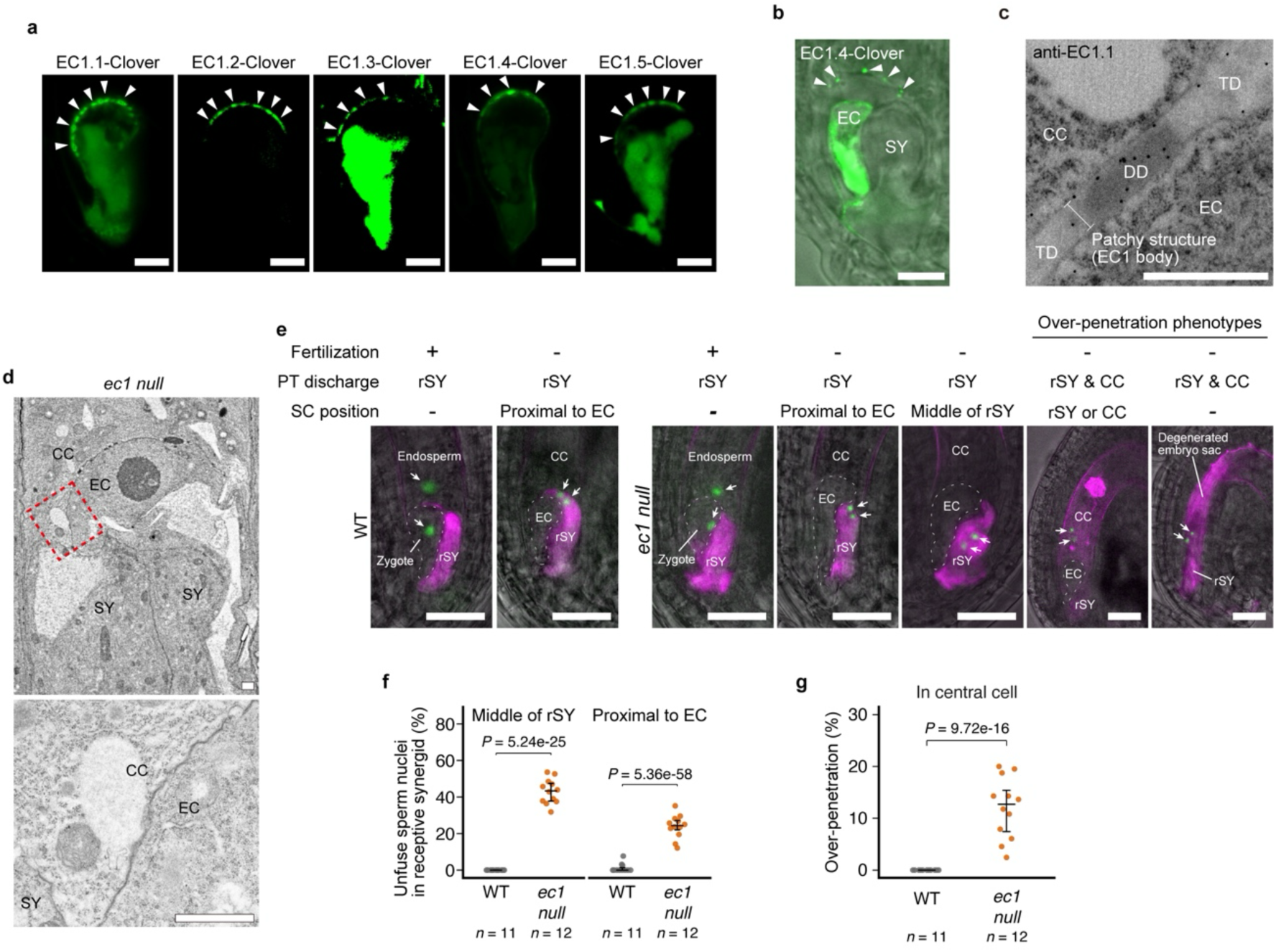
EC1 peptides form patchy extracellular structures. **a**, Localization of five *Arabidopsis* EC1 peptides in mature ovules. Arrowheads indicate extracellular punctate structures at the EC–CC interface. See also Supplementary Video 7. **b,** Plasmolysis assay of ovules expressing EC1.4-Clover. Following plasmolysis, EC1.4-Clover fluorescence remained associated with extracellular punctate structures, indicating extracellular localization. **c,** Representative immunogold electron micrograph using an anti-EC1.1 antibody (n = 3). Gold particles are predominantly associated with electron-dense domains within an extracellular patch (EC1 body). See also Extended Data Fig. 4. **d,** Representative transmission electron micrographs of the EC–CC interface in *ec1* quintuple mutant (*ec1 null*) ovules (n = 11). Extracellular patches are absent from *ec1 null* ovules. Bottom, magnified view of the dashed box shown above. **e,** Representative confocal images showing sperm-cell behavior following pollen tube discharge in wild-type (357 ovules from 11 pistils) and *ec1 null* (442 ovules from 12 pistils) ovules. After double fertilization, HTR10-Clover labels the zygote and endosperm nuclei. In *ec1 null* ovules, sperm cells frequently remain within the receptive synergid or are displaced from the EC–CC interface. Cytosolic tagRFP occasionally leaked into the central cell, and catastrophic over-penetration destroyed 23 of 54 affected ovules. Ovules were analyzed 7.5 h after pollination, immediately after pollen tube discharge, using *PTC1.0* pollen. **f,** Percentages of unfused sperm nuclei remaining within the receptive synergid. **g,** Percentages of ovules exhibiting over-penetration. In (**f**) and (**g**), dots represent individual pistils; horizontal bars indicate medians and quartiles. Sample sizes and exact *P* values from likelihood-ratio tests of binomial logistic models are shown in the figure. See also Supplementary Video 8. Abbreviations: EC, egg cell; CC, central cell; SY, synergid cell; rSY, receptive synergid; SN, sperm nucleus; DD, electron-dense domain; TD, translucent domain. Scale bars, 10 µm in (**a**), (**b**); 1 µm in (**c**) and (**d**); 20 µm in (**e**).

Plasmolysis assays showed that EC1.4-Clover fluorescence continued to label the extracellular structures rather than retracting with the egg-cell plasma membrane (Fig. 3b). Immunogold electron microscopy independently confirmed the presence of EC1 proteins within the extracellular patches (Fig. 3c). Together, these findings revise the current model of EC1 secretion and identify extracellular patches as protein assemblies enriched in EC1 peptides. We designate these extracellular structures EC1 bodies. To determine whether EC1 bodies are required for gamete pit formation, we analysed the *ec1* quintuple mutant (*ec1 null*)^24^. EC1 bodies were absent from the EC–CC interface, and localized EC–CC dissociation failed to occur during pollen tube discharge (Fig. 3d,e). Consequently, released sperm cells frequently remained within the receptive synergid or were displaced by abnormal cytoplasmic backflow. Time-lapse imaging further revealed that sperm cells initially approached the EC–CC interface but were immediately displaced toward the receptive synergid by abnormal cytoplasmic backflow following pollen tube discharge (Extended Data Fig. 5 and Supplementary Video 8). Strikingly, pollen tube contents frequently breached the central-cell plasma membrane, resulting in catastrophic over-penetration into the central-cell cytoplasm (Fig. 3f,g). These findings establish EC1 bodies as extracellular protein assemblies that organize the extracellular architecture required for controlled gamete pit formation. The proteinaceous nature of EC1 bodies raised the question of how these extracellular protein assemblies are organized.

## Amyloid-like property of EC1 body

Previous studies established the signalling activity of EC1 using synthetic peptides encompassing the conserved S1 and S2 motifs, because recombinant or synthetic full-length EC1 proteins were not available for bioactivity assays^2,25–28^. Even these synthetic peptides exhibited limited solubility owing to their hydrophobic nature, preventing accurate estimation of their effective concentration. In retrospect, these longstanding technical challenges may reflect an intrinsic propensity of EC1 proteins to self-assemble.

To examine this possibility, we first evaluated the aggregation propensity of EC1 computationally. Aggrescan3D^29^ identified aggregation-prone residues within both conserved S1 and S2 motifs, suggesting that these evolutionarily conserved regions contribute not only to signalling activity but also to intermolecular assembly (Fig. 4a,b). Furthermore, Aggrescan4D^30^ predicted that EC1 aggregation propensity increases under mildly acidic conditions, consistent with preferential EC1 assembly after secretion into the extracellular environment, which is maintained at a lower pH than the cytosol (Fig. 4c).

**Fig. 4.**
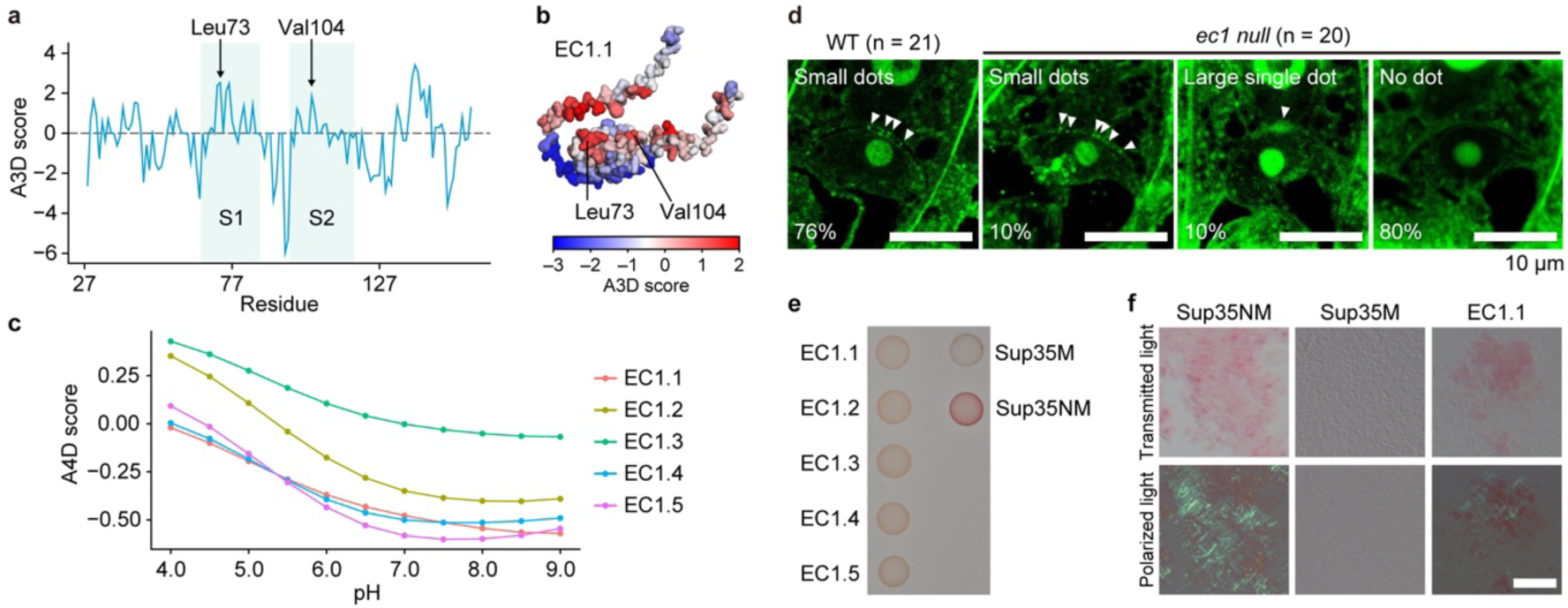
EC1 proteins exhibit aggregation-prone and amyloid-like properties. **a**, Aggrescan3D (A3D) score plot of EC1.1 after removal of the N-terminal signal sequence. Conserved S1 and S2 signature motifs are highlighted in pale blue. Similar aggregation-prone residues were predicted within the conserved S1 and S2 motifs of all five *Arabidopsis* EC1 proteins. **b,** AlphaFold2 structural model of mature EC1.1. Residues are colored according to Aggrescan3D aggregation scores. The S1 and S2 motifs form prominent aggregation-prone surfaces. **c,** Aggrescan4D prediction of mature EC1 proteins under different pH conditions. Aggregation scores increased under acidic conditions, consistent with enhanced aggregation propensity in the extracellular environment. **d,** Representative FSB staining of wild-type (n = 21) and ec1 null (n = 20) ovules. Arrowheads indicate extracellular amyloid-like deposits at the EC–CC interface. **e,** C-DAG assay showing extracellular amyloid formation by mature EC1 proteins secreted from bacteria. Yeast-derived Sup35NM and Sup35M served as positive and negative controls, respectively. **f,** Polarized-light observation of the bacterial colonies shown in (**e**). Congo Red-associated apple-green birefringence indicates extracellular amyloid formation by EC1 proteins, comparable to Sup35NM. See also Supplementary Video 9. Abbreviations: EC, egg cell; CC, central cell; SY, synergid cell. Scale bars, 10 µm.

Interestingly, the conserved core sequence of EC1 shares similarity with cereal prolamins, which assemble into electron-dense protein bodies displaying amyloid-like properties^31–34^. Because EC1 bodies likewise contain electron-dense domains enriched in EC1 proteins (Fig. 3c; Extended Data Fig. 4), we examined whether these extracellular assemblies also exhibit amyloid-like characteristics. Consistent with this possibility, independent analysis using WALTZ^35^ identified a prominent amyloidogenic region within the conserved S1 motif (Extended Data Fig. 6). FSB staining revealed strong signals in wild-type EC1 bodies but not in the *ec1 null* mutant, indicating that amyloid-like structures depend on EC1 (Fig. 4d). Correlative light and electron microscopy further demonstrated that FSB-positive signals correspond to the electron-dense domains of EC1 bodies (Extended Data Fig. 7), identifying these domains as the principal sites of amyloid-like organization.

To determine whether EC1 proteins themselves possess an intrinsic capacity to form amyloid-like assemblies, we employed the curli-dependent amyloid generator (C-DAG) system^36^, using the authentic yeast prion protein Sup35NM and the non-amyloidogenic Sup35M as positive and negative controls, respectively. Extracellular expression of EC1 proteins produced Congo Red-positive colonies with characteristic apple-green birefringence under polarized light, a hallmark of amyloid structures, comparable to Sup35NM but absent from Sup35M controls (Fig. 4e,f, Supplementary Video 9).

Collectively, these findings indicate that EC1 proteins possess intrinsic self-assembly properties and a subset of which forms amyloid-like assemblies within EC1 bodies, providing a plausible molecular basis for the formation and stabilization of extracellular EC1 bodies. These observations raised the question of whether the structural properties of extracellular protein assemblies themselves contribute to EC1 body function.

## Partial functional substitution of EC1 bodies by heterologous extracellular protein assemblies

To determine whether the biological function of EC1 bodies depends on the molecular identity of EC1 or on the structural properties of extracellular protein assemblies, we generated transgenic *ec1 null* plants carrying a Sup35NM replacement construct, in which the yeast-derived amyloidogenic prion domain Sup35NM was redirected into the egg-cell secretory pathway by fusion to the EC1.1 signal peptide and tagged with mNeonGreen to visualize its localization (Fig. 5a). As a non-amyloidogenic control, we generated an equivalent Sup35M replacement construct lacking the N-terminal prion-forming domain. The resulting transgenic plants are hereafter referred to as the *Sup35NM replacement* (*Sup35NM repl*) and *Sup35M replacement* (*Sup35M repl*) lines, respectively. Both fusion proteins accumulated at the EC–CC interface, although Sup35NM-mNeonGreen consistently formed larger extracellular deposits than Sup35M-mNeonGreen (Fig. 5a).

**Fig. 5.**
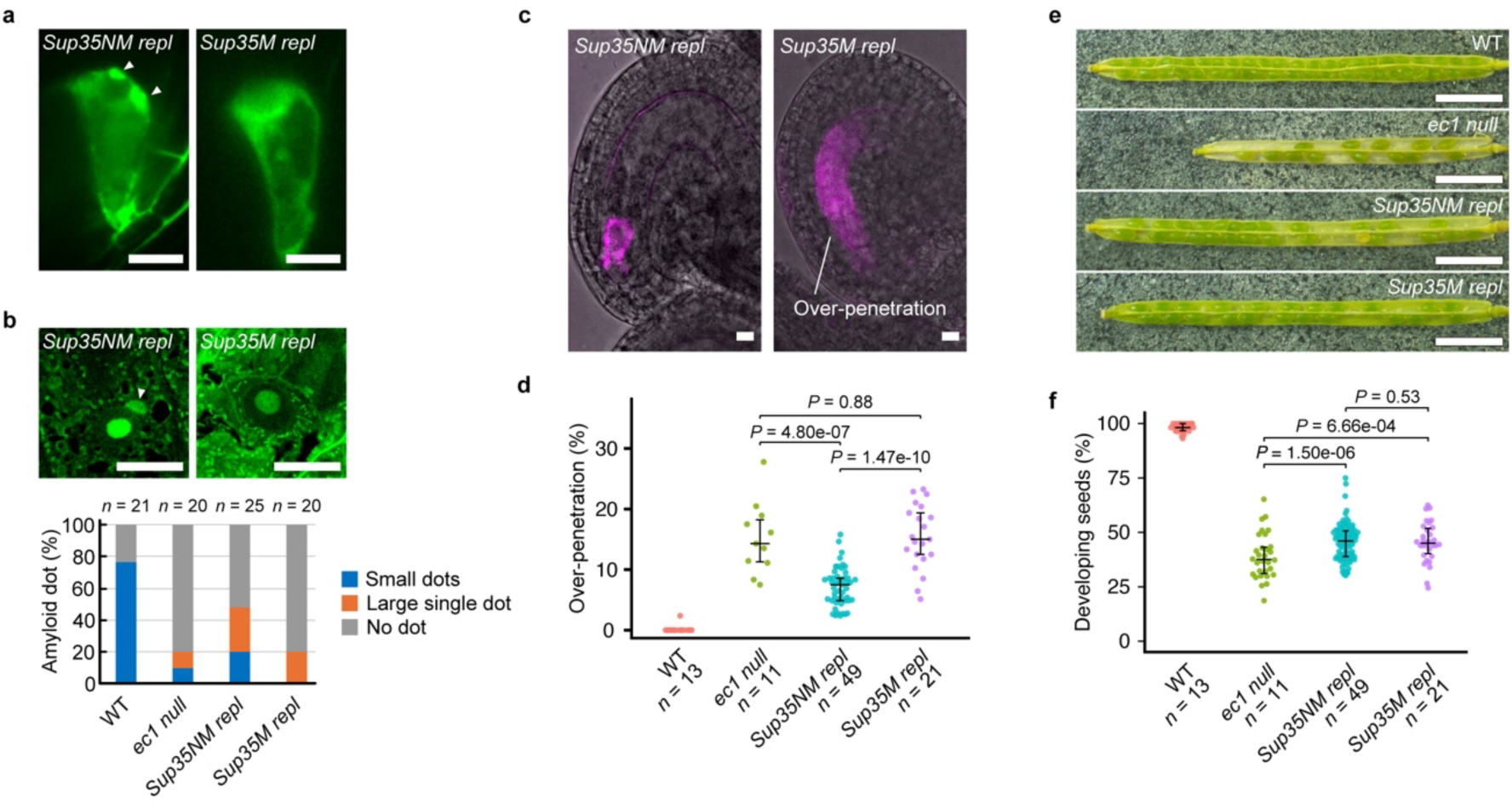
Partial functional substitution of EC1 bodies by a heterologous extracellular protein assembly. **a**, Representative confocal images of mature ovules in the *Sup35NM replacement* (*Sup35NM repl*) and *Sup35M replacement* (*Sup35M repl*) plants. In these plants, the amyloidogenic yeast prion domain Sup35NM or the non-amyloidogenic Sup35M control, fused to the EC1.1 signal peptide and mNeonGreen, were expressed from the *EC1.1* promoter in the *ec1 null* background. Large extracellular deposits (arrowheads) were frequently observed in *Sup35NM repl* ovules (29.9%, n = 107), but rarely in *Sup35M repl* ovules (0.8%, n = 133). **b,** Representative FSB staining of the *Sup35NM repl* and *Sup35M repl* ovules. Lower panels summarize the frequencies of ovules exhibiting small dots, large single dot, or no detectable FSB-positive extracellular deposits. Quantification of wild-type and *ec1 null* ovules is reproduced from Fig. 4d for comparison. **c,** Representative confocal images showing pollen tube discharge in *Sup35NM repl* and *Sup35M repl* ovules. **d,** Percentages of ovules exhibiting over-penetration after pollen tube reception. Ovules were analyzed 7.5 h after pollination, immediately after pollen tube discharge, using PTC1.0 pollen. **e,** Representative siliques of wild-type, *ec1 null*, *Sup35NM repl*, and *Sup35M repl* plants. **f,** Seed set of the indicated genotypes. In (**d**) and (**f**), dots represent individual pistils and siliques, respectively; horizontal bars indicate medians and quartiles. Sample sizes and exact Benjamini–Hochberg-adjusted *P* values from pairwise comparisons of binomial logistic models are shown in the figure. In *Sup35NM repl,* data from three independently generated lines were combined. See also Extended Data Fig. 9 for the results of individual lines. Abbreviations: EC, egg cell; CC, central cell; SY, synergid cell; rSY, receptive synergid. Scale bars, 10 µm in (**a–c**); 2 mm in (**e**).

We next examined whether these heterologous extracellular deposits acquired amyloid-like organization. FSB staining revealed that most *ec1 null* ovules lacked detectable extracellular amyloid-like deposits, although small residual puncta or occasional larger deposits remained in a minority of ovules (Fig. 4d, Fig. 5b). By contrast, *Sup35NM repl* lines substantially increased the frequency of FSB-positive extracellular assemblies and frequently formed enlarged extracellular FSB-positive deposits compared with the punctate structures observed in wild-type ovules. *Sup35M repl* lines occasionally produced larger FSB-positive deposits but did not increase the overall frequency of FSB-positive structures. Transmission electron microscopy further revealed prominent electron-dense extracellular domains within Sup35NM-derived extracellular assemblies that resembled the dense domains of native EC1 bodies (Fig. 3c, Fig. 5b; Extended Data Fig. 8). Thus, redirecting an unrelated amyloidogenic protein into the egg-cell secretory pathway was sufficient to reconstruct extracellular protein assemblies sharing key structural features of native EC1 bodies.

We next asked whether these heterologous extracellular protein assemblies could substitute for EC1 body function. Whereas pollen tube discharge frequently resulted in catastrophic over-penetration of pollen tube contents into the central-cell cytoplasm in *ec1 null* ovules, this phenotype was markedly suppressed in *Sup35NM repl* lines but only modestly affected in *Sup35M repl* lines (Fig. 5c,d; Extended Data Fig. 9). Thus, amyloidogenic extracellular protein assembly, rather than extracellular protein deposition alone, substantially contributed to restricting pollen tube contents at the EC–CC interface. Consistent with this improvement, female fertility was partially restored in *Sup35NM repl* lines, whereas *Sup35M repl* lines exhibited only a modest increase in seed set (Fig. 5e,f; Extended Data Fig. 9). Together, these findings demonstrate that heterologous extracellular protein assemblies partially substitute for EC1 body function. The incomplete complementation likely reflects structural differences between Sup35-derived assemblies and native EC1 bodies, indicating that extracellular protein architecture is a major functional determinant of the controlled confinement of pollen tube contents, whereas additional architectural and molecular features optimize native EC1 body function.

## Discussion

Since the discovery of double fertilization more than a century ago^37^, how immotile sperm cells are accurately delivered to two female gametes immediately after pollen tube discharge has remained one of the central unresolved questions in flowering plant reproduction^3,4,38^. Our findings show that successful sperm-cell reception depends on the rapid construction of a transient extracellular fertilization niche generated by localized dissociation of the EC–CC interface. This process is orchestrated by EC1 bodies, extracellular protein assemblies that transiently remodel the gamete interface and thereby constrain the path of pollen tube contents toward the fertilization site. Rather than representing a passive consequence of explosive pollen tube discharge, gamete pit formation emerges as an actively organized cellular event that ensures precise sperm-cell delivery while preventing catastrophic invasion of the central-cell cytoplasm (Extended Data Fig. 10).

Our findings also fundamentally revise the current view of EC1 function. EC1 peptides were originally identified as gamete-activating signals that induce redistribution of the sperm membrane fusion factor GCS1/HAP2 immediately before gamete fusion^2^. We now show that EC1 peptides are secreted before pollen tube discharge and accumulate extracellularly as a major component of EC1 bodies (Fig. 3a–d; Extended Data Fig. 4), where they promote the assembly of a transient extracellular structure required for accurate sperm-cell delivery. Our live-imaging analyses further indicate that the primary defect in *ec1 null* mutants arises immediately after pollen tube discharge, suggesting that previously reported abnormalities in sperm-cell positioning^25^ likely represent downstream consequences of defective initial sperm-cell reception rather than independent defects in gamete adhesion. These observations suggest that accurate sperm-cell positioning is established before EC1-dependent activation of gamete fusion, thereby redefining the temporal sequence of EC1 function during fertilization. Overall, EC1 performs two temporally distinct functions during fertilization: first as a structural organizer that establishes the extracellular protein architecture of the gamete interface, and subsequently as a signalling peptide that activates sperm cells for membrane fusion (Extended Data Fig. 10). The evolutionary conservation of the aggregation-prone S1 and S2 motifs further raises the possibility that this structural role has been maintained together with the well-established signalling activity of EC1 throughout angiosperm evolution.

More broadly, our findings uncover a cell-wall-independent mode of cell-interface remodeling mediated by extracellular protein assemblies. Cell separation in plants has generally been understood as a consequence of enzymatic modification or degradation of the cell wall^39–42^. By contrast, EC1 bodies transiently remodel the geometry of the EC–CC interface through extracellular protein assembly without extensive cell-wall remodeling. In the context of flowering plant fertilization, such a mechanism may be particularly advantageous because it rapidly creates a permissive extracellular path for sperm-cell delivery while preserving the mechanical integrity of both female gametes immediately before membrane fusion. Unlike irreversible cell-wall degradation, transient protein-mediated remodeling allows the interface to be established with high spatial precision and to disappear immediately after fertilization. Our replacement experiments further support that this function depends not solely on the molecular identity of EC1, but also on the extracellular protein architecture that shapes cell-interface geometry (Fig. 5). Together, these observations suggest that extracellular protein architecture provides a previously unrecognized mechanism for transient cell-interface remodeling during highly coordinated cellular interactions.

Finally, our findings further raise the possibility that extracellular protein architecture represents a broader biological strategy for generating dynamic cellular microenvironments. Although demonstrated here in flowering plant fertilization, transient extracellular protein assemblies could provide a versatile means of locally organizing intercellular geometry whenever cellular interactions require both rapid assembly and rapid dissolution. Future studies will determine whether analogous protein-mediated remodeling mechanisms contribute to other developmental or reproductive processes. By identifying extracellular protein architecture as a structural principle underlying dynamic cell-interface remodeling, our work expands the conceptual framework through which extracellular proteins are understood in cell–cell communication.

## Methods

### Plant material and growth conditions

Columbia-0 (Col-0) was used as the wild-type strain. Seeds were surface sterilized with sterilization solution (0.1% Tween 20, 2% Plant Preservative Mixture (Plant Cell Technology, Washington DC, US)) and incubated at 4°C for several days and sown on Murashige and Skoog (MS) medium containing 1% sucrose and appropriate antibiotics. Approximately 10 days after germination, plants were transferred to soil. Plants were grown at 22°C under continuous light or standard long day conditions.

### Plasmids and transgenic plants

The *pHTR10:HTR10-Clover* marker line^43^ and the *pMYB98:GFP-PIP2a* marker line^44^ were described previously. The pSAN37, a destination vector carrying *DD65* promoter, was reported previously^44^. The *pLAT52:tagRFP* plasmid was a gift from S. Nishikawa (Niigata Univ.). The MU2000, a pGWB501^45^ containing the *EC1.1* promoter, was provided by M. Ueda (Tohoku Univ.). The pVS72 and pVS105, vectors for bacterial expression of Sup35NM and Sup35M, were purchased as the C-DAG Amyloidogenicity Kit^36^ *via* kerafast website (https://www.kerafast.com). The *pLAT52:tagRFP* pollen tube cytosolic marker gene was introduced into the *pHTR10:HTR10-Clover* sperm nuclear marker plant to produce the *PTC1.0* plants. Generation of *Arabidopsis* transformants was performed by floral dipping method^46^ using an *Agrobacterium tumefaciens* strain GV3101. Plasmid construction procedures are described in Supplementary Methods.

### Observation of mature ovules

Emasculated pistils or pollinated pistils were harvested, and the carpel walls were removed on a glass slide. Then, the pistils were split into half and mounted on 5% sucrose, 0.3M sorbitol solution, or an ovule culture medium containing Nitsch basal salt and 5% trehalose dihydrate^47^. Confocal images were obtained by Leica SP8 TCS (Leica, Wetzlar, Germany) or IX73 inverted microscope (Evident, Tokyo, Japan) equipped with a spinning disk confocal scanning unit (CSU-W1; Yokogawa, Tokyo, Japan) and an sCMOS camera (Zyla 4.2; Andor, Belfast, Northern Ireland).

### FSB staining

Pistils were harvested one day after emasculation, and the carpel walls and stigmas were removed. Samples were fixed in a PFA/GA solution (4% paraformaldehyde and 2% glutaraldehyde in 0.05 M phosphate buffer) via vacuum infiltration for 30 min (15 min×2) and incubated at 4°C for at least 24 h. Fixed samples were dehydrated through a graded ethanol series (70–100%) and infiltrated with Technovit 7100 resin (Heraeus Kulzer, Hanau, Germany). The ovules were then dissected and embedded in Technovit 7100 using beam capsules (Nisshin EM, Tokyo, Japan). For block mounting, Technovit 3040 was applied. Longitudinal sections (1 μm thick) were prepared using a microtome (RM2245, Leica, Wetzlar, Germany) and mounted on glass slides. For the detection of amyloid aggregates, sections were stained with 0.001% (w/v) FSB (1-fluoro-2,5-bis(3-carboxy-4-hydroxystyryl)benzene) in 50% ethanol for 30 min. After rinsing with water and drying, fluorescence was observed using a confocal laser scanning microscope (SP8 TCS, Leica) equipped with a 63× glycerol immersion objective and a hybrid detector (HyD). FSB fluorescence was excited at 405 nm and detected within a 436–560 nm emission range.

For the correlative light and electron microscopy (CLEM), Technovit-embedded samples were sectioned at a thickness of 500 nm using an EM UC7 ultramicrotome (Leica, Wetzlar, Germany). The sections were initially stained with FSB, and those containing ovules with extracellular amyloid aggregates were identified *via* confocal microscopy. These target sections were then stained with 0.4% uranyl acetate for 10 min, rinsed with water, and dried. Subsequently, the sections were treated with a lead staining solution (Sigma-Aldrich, Japan) for 1 min and subjected to osmium coating. Ultrastructural observations were performed using an SU8220 field-emission scanning electron microscope (FE-SEM) equipped with the Mirror CLEM system (Hitachi High-Tech, Tokyo, Japan). FE-SEM images were acquired with an yttrium aluminum garnet (YAG) backscattered electron detector at an acceleration voltage of 5 kV. The morphology of extracellular patches was characterized by correlating the FE-SEM images with the pre-acquired FSB fluorescence data.

### Transmission electron microscopy (TEM)

Emasculated pistils were fixed in a solution containing 4% paraformaldehyde, 2% glutaraldehyde, and 50 mM sodium cacodylate at pH7.4 for several days at 4°C. The samples were washed in buffer and post-fixed for 6 h in 2% aqueous osmium tetroxide at 4°C. The specimens were then dehydrated in a graded ethanol series, transferred into propylene oxide, infiltrated, and embedded in Quetol 651. Series of thin-sections (80 nm) were stained with 2% aqueous uranyl acetate and lead citrate, and examined at 80 kV under a JEOL JEM 1400 Plus electron microscope (JEOL Ltd.). Digital images were taken with a CCD camera (VELETA; Olympus Soft Imaging Solutions).

### Immunogold electron microscopy

For SBT4.13-Clover, unfertilized *Arabidopsis* ovules (1 d after emasculation) were processed for immunogold electron microscopy as follows. Samples were fixed in 4% (w/v) paraformaldehyde and 0.1% (v/v) glutaraldehyde in 0.1 M phosphate buffer (pH 7.4) at 4 °C for 90 min, washed three times in 0.1 M phosphate buffer (15 min each), dehydrated in graded ethanol (50% and 70%) at 4 °C for 30 min each, infiltrated with ethanol:LR White (50:50, 30 min ×3) followed by 100% LR White (4 °C, 30 min ×3), and UV-polymerized at 4 °C overnight. Ultrathin sections (90 nm; Ultracut UCT; Leica) were collected on nickel grids and labeled with rabbit anti-GFP polyclonal antibody in 1% BSA/PBS for 2 h at room temperature, followed by 10-nm gold-conjugated goat anti-rabbit IgG for 1 h at room temperature. After post-fixation in 2% (v/v) glutaraldehyde in 0.1 M phosphate buffer, sections were counterstained with 2% uranyl acetate (15 min) and lead stain solution (3 min) and imaged on a transmission electron microscope (JEM-1400Plus, JEOL).

For EC1.1 immunogold labeling, unfertilized Arabidopsis ovules (1 d after emasculation) were fixed in 4% (w/v) paraformaldehyde and 2% (v/v) glutaraldehyde in 0.05 M phosphate buffer (pH 7.4) at 4 °C overnight. Samples were washed in 0.05 M phosphate buffer (5 min, six times) and dehydrated through a graded methanol series (25%, 50%, 75%, 90% and 100%; two changes each, 30 min per change). Samples were infiltrated with LR White resin (50% and 100%; each for 0.5 d), embedded in gelatin capsules, and polymerized by UV irradiation at −20 °C for 72 h. Ultrathin sections (70 nm) were cut using an ultramicrotome (EM UC7, Leica) and collected on nickel grids. A rabbit polyclonal anti-AtEC1.1 antibody was generated commercially (Cosmo Bio) using a synthetic peptide corresponding to AtEC1.1 residues 42–54 (TSLVYRLKLDEDT) as the immunogen and affinity-purified on a peptide column. For immunogold labeling, grids were blocked with Block Ace (KAC, Japan) and incubated for 1 h with rabbit pre-immune serum (1:100) or rabbit anti-AtEC1.1 antibody (1:100). After washing with 1:10-diluted Block Ace containing 0.02% Triton X-100, grids were incubated for 30 min with 18-nm gold-conjugated anti-rabbit IgG (1:30). Grids were electron stained with 4% uranyl acetate for 10 min before imaging on a transmission electron microscope (JEM 1400 Flash, JEOL).

### FIB-SEM

The FIB–SEM analysis were performed essentially as described previously^48–50^. Unfertilized *Arabidopsis* ovules (1 d after emasculation) were pre-fixed with 4% paraformaldehyde and 2% glutaraldehyde in 0.05 M cacodylate buffer (pH 7.4) at 4 °C overnight. The samples were post-fixed with 2% osmium tetroxide in the same buffer at 4 °C for 1 h. After being thoroughly rinsed, samples were treated with 1% thiocarbohydrazide aqua at room temperature for 20 min, then rinsed well, and post-fixed again with 2% osmium tetroxide aqua at room temperature for 1 h. After being thoroughly rinsed, samples were stained with 1% uranyl acetate aqua at 4 °C overnight, then rinsed, and stained with 0.03 M lead aspartate aqua at 60 °C for 30 min. Samples were dehydrated in graded ethanol series (50%, 70%, 90% and 100%), transferred into propylene oxide, infiltrated, and embedded in resin (Quetol 651, Nissin-EM). Resin blocks were trimmed, carbon-coated, and cut and observed repeatedly using FIB-SEM (MI-4000L, Hitachi High-Tech). FIB milling (cutting) conditions were as follows: accelerating voltage, 30 kV; beam current for milling to prepare or to cut, 1.6 or 1.2 nA, respectively; cutting interval (z-step), 25 nm. SEM (image capturing) conditions were as follows: accelerating voltage, 1.0 kV; beam current, 150 pA; working distance, 2.0 mm; detector, Upper (in-lens for secondary electron) + EsB (energy and angle selective backscattered electron); image size, 2000 × 2000 pixels; field of view, 50 × 50 µm; pixel size, 25 nm per pixel; color depth, 8-bit (256 gray scales).

The images of serial sections were processed using software Fiji^51^; gray scale was inverted, brightness and contrast were adjusted, and then the serial images were aligned using the plugin “Register Virtual Stack Slices” (Translation -no deformation). On the processed serial images (approx. 400 sections, totally 10 µm z-thickness), every 6 images were extracted (z-step: 25 nm × 5 sections = 125 nm interval), and contours of the cell walls were traced manually, and the segmented area of two synergid cells, a central cell, an egg cell, and the gap between the walls of central and egg cells were filled with monotone colors using software Affinity Designer (Ver. 1.10.8, Serif Europe Ltd.). The segmented images were used for 3D reconstruction using software Avizo (ver. 9.0, Thermo Fisher Scientific Inc.).

### Amyloidogenicity assay

Amyloidogenicity of Arabidopsis EC1 peptides (EC1.1‒EC1.5) was assessed using the curli-dependent amyloid generator (C-DAG) system (C-DAG Amyloidogenicity Kit; Kerafast) following the published protocol^36^ and the manufacturer’ s instructions. Coding sequences lacking the signal peptide were cloned into the pExport vector as N-terminal fusions to the CsgA signal sequence (CsgAss‒EC1.1‒EC1.5) with a C-terminal His_6_ tag, using NotI and XbaI restriction sites. Exporter cells (E. coli VS45) were transformed by heat shock and selected on LB agar containing carbenicillin (100 µg ml^−1^) and chloramphenicol (25 µg ml^−1^). For Congo red colony-color and birefringence assays, transformants were spotted onto Congo red inducing plates (LB agar supplemented with 100 µg ml^−1^ carbenicillin, 25 µg ml^−1^ chloramphenicol, 0.2% (w/v) L-arabinose, 1 mM IPTG, and 10 µg ml^−1^ Congo red) and incubated at 22 ℃ for 5 days. For birefringence, bacterial spots were resuspended on the CR inducing plate with PBS, and 5 µl of the suspension was placed on a glass slide, air-dried for 2 min, coverslipped, and examined by transmitted-light microscopy between crossed polarizers (IX71; Olympus, Japan; FLOYD; Wraymer, Japan). Kit-provided positive and negative control plasmids (pVS72 and pVS105, encoded Sup35NM and Sup35M respectively) were processed in parallel under identical conditions.

### Plasmolysis analysis

Unfertilized ovules from the *pEC1.4:EC1.4‒Clover* were obtained by dissecting stage 12c flowers at 1 d after emasculation. Dissected ovules were incubated in a hyperosmotic enzyme solution containing 1% cellulase (Worthington Biochemical Corporation, USA), 0.3% macerozyme R-10 (Yakult, Japan) and 0.05% pectolyase (Kyowa Kasei, Japan) (pH 7.0), modified from the egg-cell isolation enzyme solution described previously by substituting the osmoticum with 1 M sorbitol. After incubation for 30 min at 24 °C, ovules were observed with a spinning disk confocal scanning unit (CSU-W1; Yokogawa, Japan) and an sCMOS camera (Zyla 4.2; Andor, Northern Ireland).

### Semi-*in vivo* fertilization assay

For the time-lapse imaging of pollen tube discharge, we modified the protocol of semi-*in vivo* fertilization assay described by Hamamura et al1. In brief, thin-layered pollen-tube growth medium (14% sucrose, 0.001% boric acid, 1.27 mM Ca(NO_3_)_2_, 0.4 mM MgSO_4_, adjusted pH to 7.0 with 1 N KOH, 1.5% NuSieve GTG agarose, and 10 µM epibrassinolide) was prepared on a glass bottom dish (D11130H, Matsunami Glass IND., LTD., Japan). Stigmas from emasculated wild-type pistils were cut with a needle and placed on the medium, followed by hand-pollination with pollen from transgenic plants. After 3–5 h of incubation, pollen tubes that emerged from the cut pistils were observed by the TCS SP8 microscope.

### Aggregation assessment

Amyloid prediction was performed by using Waltz (https://waltz.switchlab.org)^35^. Signal peptides were predicted using SignalP 6.0^52^ in eukaryotic mode with slow (high-accuracy) settings. Signal peptide cleavage sites were extracted from the prediction results, and mature peptide sequences were generated by removing the signal peptide region using a custom Python script. Protein three-dimensional structures were predicted using the AlphaFold2 (ptm model) implemented in ColabFold^53^ v1.6.1, with 3 recycles and 5 model outputs. Aggregation propensity was evaluated using Aggrescan3D^29^ on the predicted structures. The relationship between structure and aggregation scores was visualized using py3Dmol (v2.5.4; https://github.com/avirshup/py3dmol), a Python interface to 3Dmol.js^54^. Each residue was colored according to its aggregation score using a gradient scale and displayed as a three-dimensional model. pH-dependent aggregation propensity was evaluated using Aggrescan4D^30^ (https://biocomp.chem.uw.edu.pl/a4d/). The resulting data were summarized and visualized using the R package ggplot2^55^.

### Statistical analysis

Statistical analyses were performed using R (version 4.4.2). For ovule phenotypes, the numbers of ovules with and without each phenotype were recorded for each pistil and analysed using binomial generalized linear models with a logit link, with genotype as the explanatory variable. Seed set was analysed similarly using the numbers of developing and non-developing seeds recorded for each silique. Percentages are shown for visualization, whereas count data were used for statistical analyses. For two-group comparisons, genotype effects were assessed using likelihood-ratio tests comparing models with and without genotype. For comparisons among multiple genotypes, pairwise contrasts of estimated marginal means were evaluated using two-sided Wald tests, with *P* values adjusted using the Benjamini–Hochberg method^56^. Sample sizes and exact *P* values, adjusted for multiple comparisons where applicable, are shown in the figures.

### Software

Images were processed by FIJI (http://fiji.sc/Fiji)^51^. Image-Pro Premier 3D (ver. 9.3, Media Cybernetics, USA).

## Data availability

The data that support the findings of this study are available from the corresponding authors upon reasonable request.

## Acknowledgements

We acknowledge M. Tsukatani and H. Ikeda for assistance in preparing materials; Y. Mizuta (Nagoya Univ.), S. Nishikawa (Niigata Univ.), T. Igawa (Chiba Univ.), T. Kawashima (Kentucky Univ.) and T. Nakagawa (Shimane Univ.) for providing plasmids or seeds; T. Higashiyama (Univ. of Tokyo), M. Y. Hirai (Riken) for continuous support for the project, Tokai-EM Inc. for immunogold electron microscopy and transmission electron microscopy. FIB-SEM observation was supported by the Nagoya University microstructural characterization platform as a program of “Nanotechnology Platform” of the Ministry of Education, Culture, Sports, Science and Technology (MEXT), Japan, and we especially thank S. Arai and S. Enomoto (Nagoya Univ.) for technical support. This work was partially supported by Japan Society for the Promotion of Science (JSPS) KAKENHI [Grant Numbers: JP17H05846, JP20H05422, JP19H04869, JP20H03280, JP20H05778, JP20H05781, JP23K17375, JP2502300 to D.M.; JP18J01963, JP19K16172, JP22K15145, JP23H04749, JP25K09678, and JP25H01828 to D.S.; JP22H04926 for “Advanced Bioimaging Support”], by the grant for 2019–2026 Research Development Fund of Yokohama City University (to D.M.), the grant for 2019–2020 Strategic Research Promotion (Nos. SK1903) of Yokohama City University (to D.M.), Toyoaki Scholarship Foundation (to D.M.). This manuscript was prepared with the assistance of ChatGPT and Gemini to improve linguistic readability. The authors have reviewed, revised and approved the final content.

## Author contributions

D.M. conceptualized and directed this project including interpretation of the experimental results, with contributions from other authors; D.S., H.O., T.K., A.Y., and K.Tonosaki conducted live-imaging, FSB staining, genetic analyses, and data analyses; D.S., S.N., T.H., and N.S. generated plasmids and plant materials; H.O., Y.G., M.S., and K.Toyooka performed CLEM and immunogold staining; H.K. analyzed amyloidogenicity of EC1 peptides using C-DAG kit; T.O. obtained FIB-SEM images and generated 3D data; D.M. and D.S. wrote the manuscript, and all authors contributed to edit the manuscript and approved the manuscript submission.

## Extended Data Figures

**Extended Data Fig. 1.**
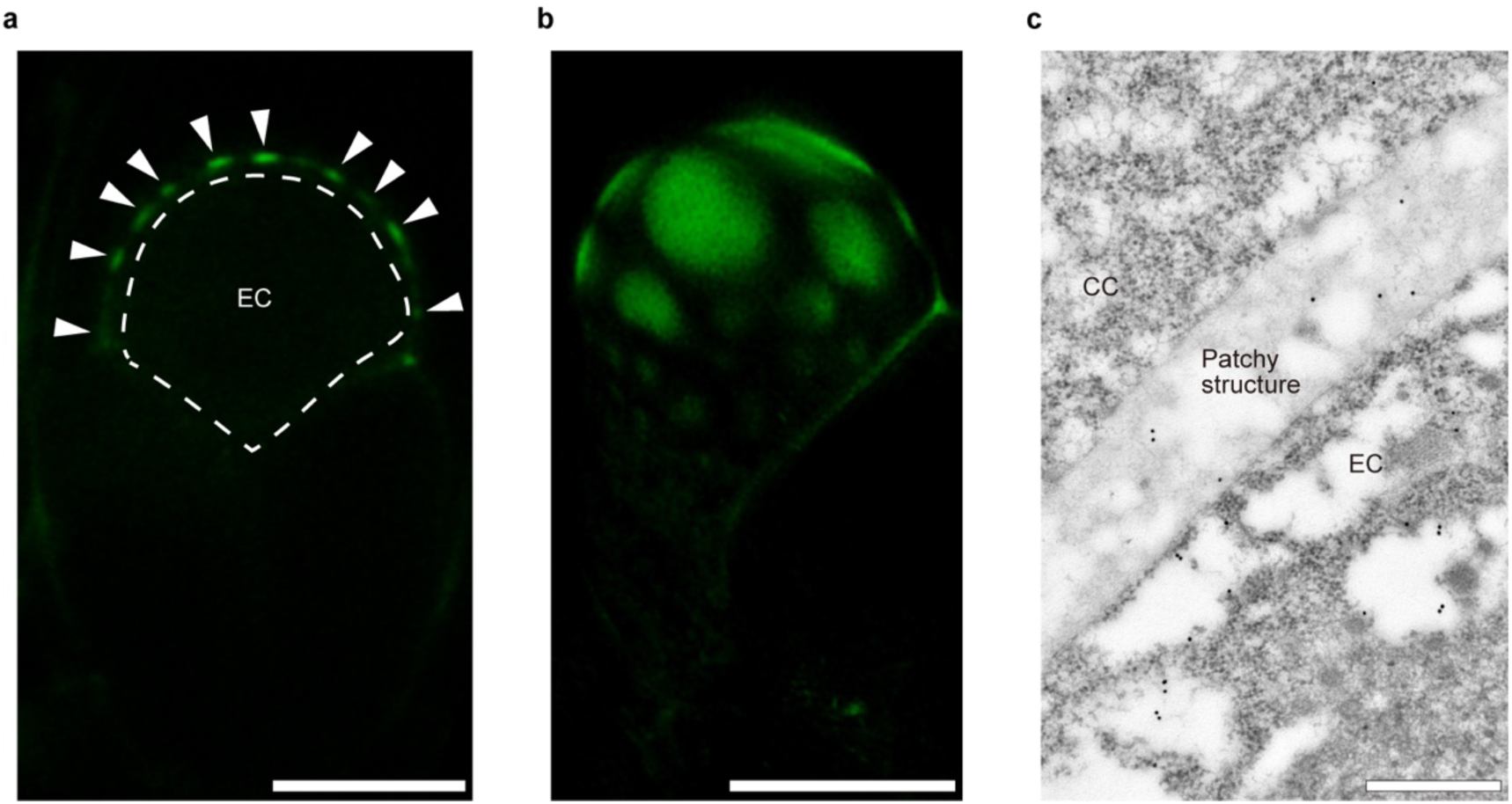
SBT4.13-Clover labels patchy extracellular structures. **a**, Patchy extracellular structures (arrowheads) in a mature ovule expressing *pSBT4.13:SBT4.13-Clover.* **b,** Three-dimensional reconstruction of the patchy extracellular structures shown in (**a**). See also Supplementary Video 5. **c,** Representative immunogold electron micrograph of the patchy extracellular structures in *pSBT4.13:SBT4.13-Clover* ovules using an anti-GFP antibody (n = 4). Abbreviations: EC, egg cell; CC, central cell. Scale bars, 10 µm in (**a**) and (**b**); 500 nm in (**c**).

**Extended Data Fig. 2.**
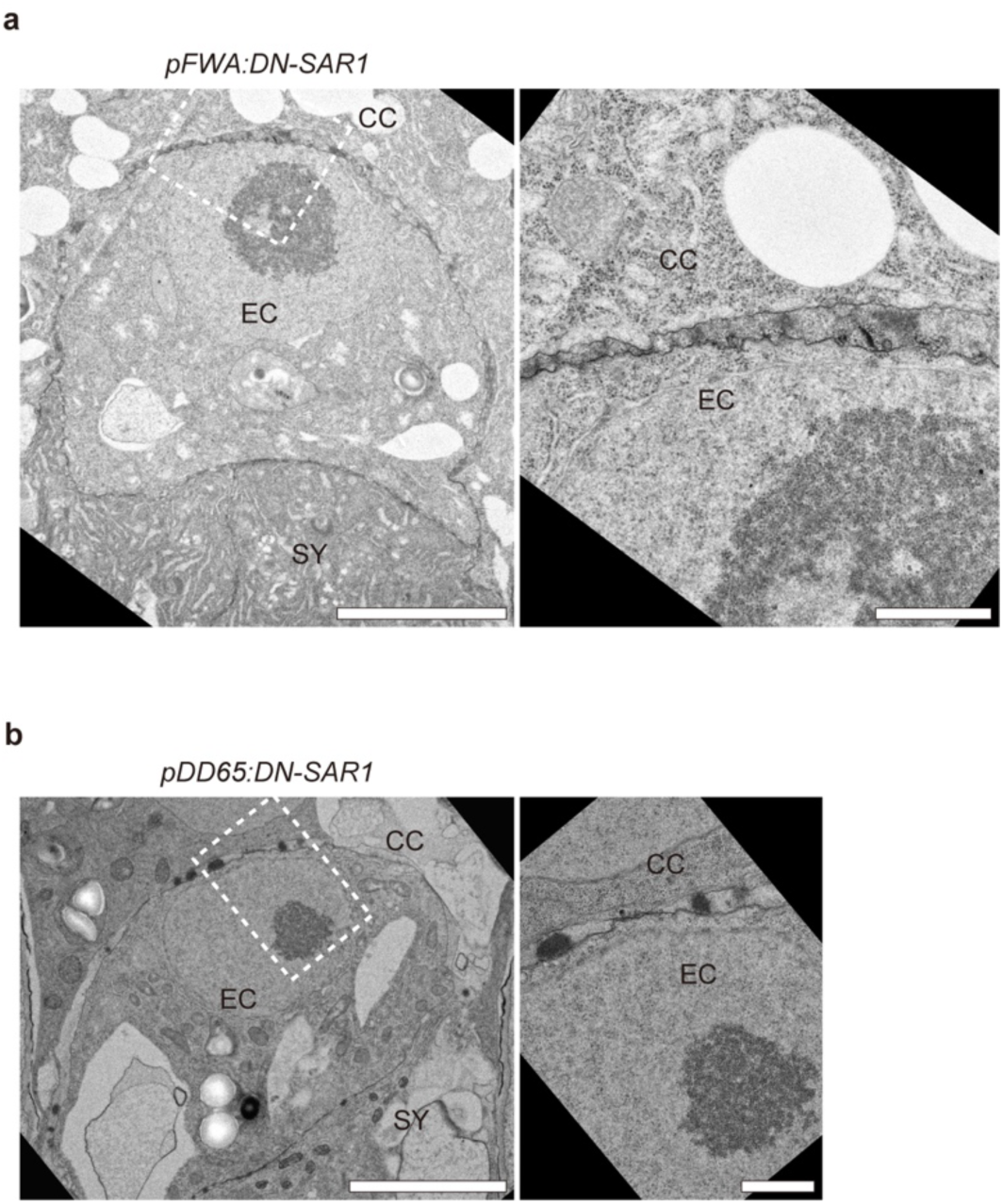
Central cell-specific DN-SAR1 expression. **a,b,** Representative transmission electron micrographs of the EC–CC interface in transgenic plants expressing DN-SAR1 under the control of the central cell-specific promoter *pFWA* (**a,** n = 5) or *pDD65* (**b,** n = 10). Right panels show magnified views of the dashed boxes in the left panels. Abbreviations: EC, egg cell; CC, central cell; SY, synergid cell. Scale bars, 5 µm in left panels; 1 µm in right panels.

**Extended Data Fig. 3.**
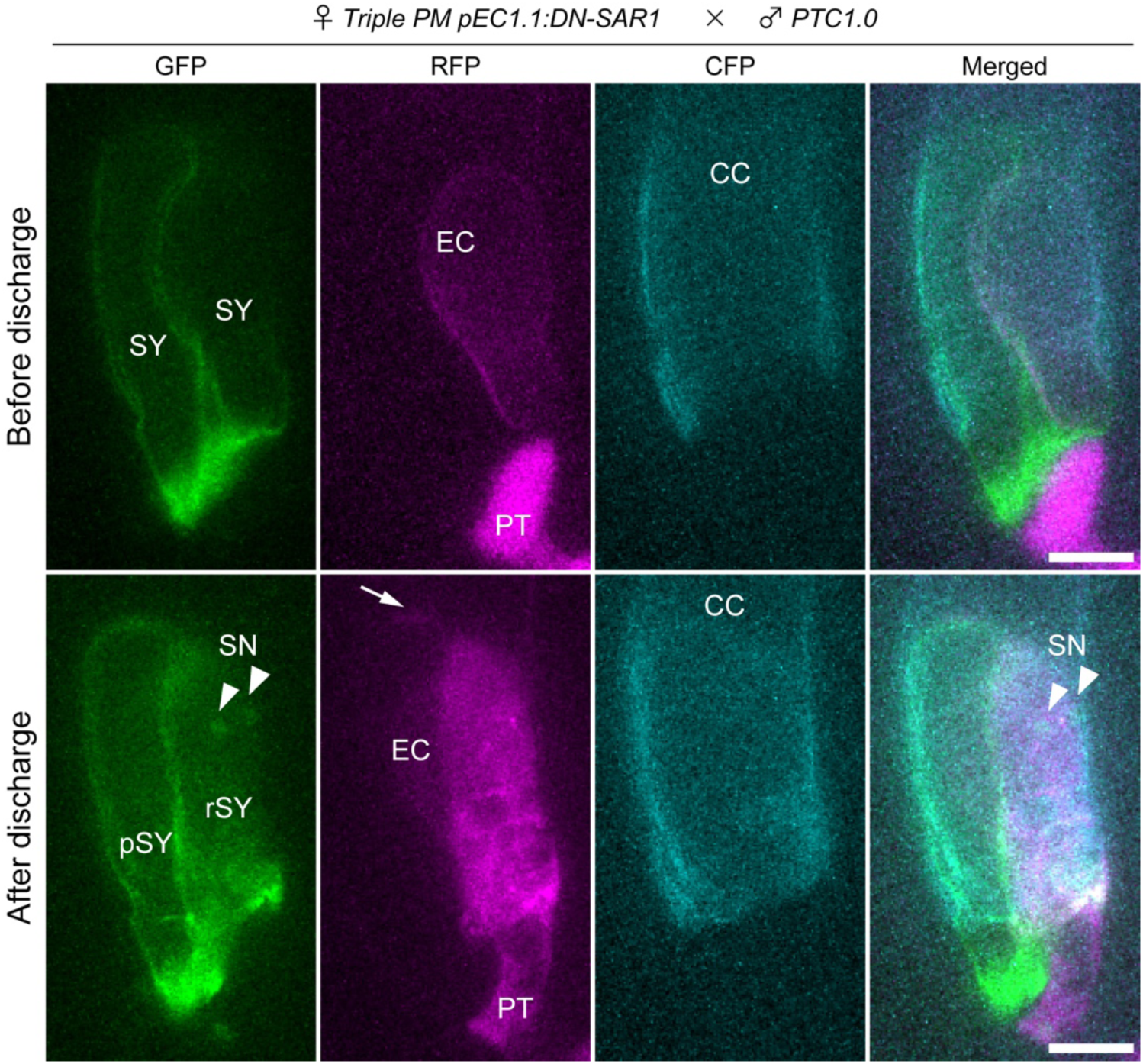
Abnormal pollen tube discharge in ovules with impaired COPII-mediated protein secretion in the egg cell. Semi-*in vivo* fertilization assay using ovules from the *pEC1.1:DN-SAR1* transgenic line carrying the *Triple PM* marker and pollen from the *PTC1.0* marker line. Maximum-intensity projections of confocal images acquired before (upper panels) and after (lower panels) pollen tube discharge are shown. Arrowheads indicate sperm nuclei. Arrow indicates slight leakage of pollen tube-derived cytosolic tagRFP. See also Supplementary Video 6. Abbreviations: SC, sperm cell; SN, sperm nucleus; PT, pollen tube; EC, egg cell; CC, central cell; SY, synergid cell; rSY, receptive synergid. Scale bars, 10 µm.

**Extended Data Fig. 4.**
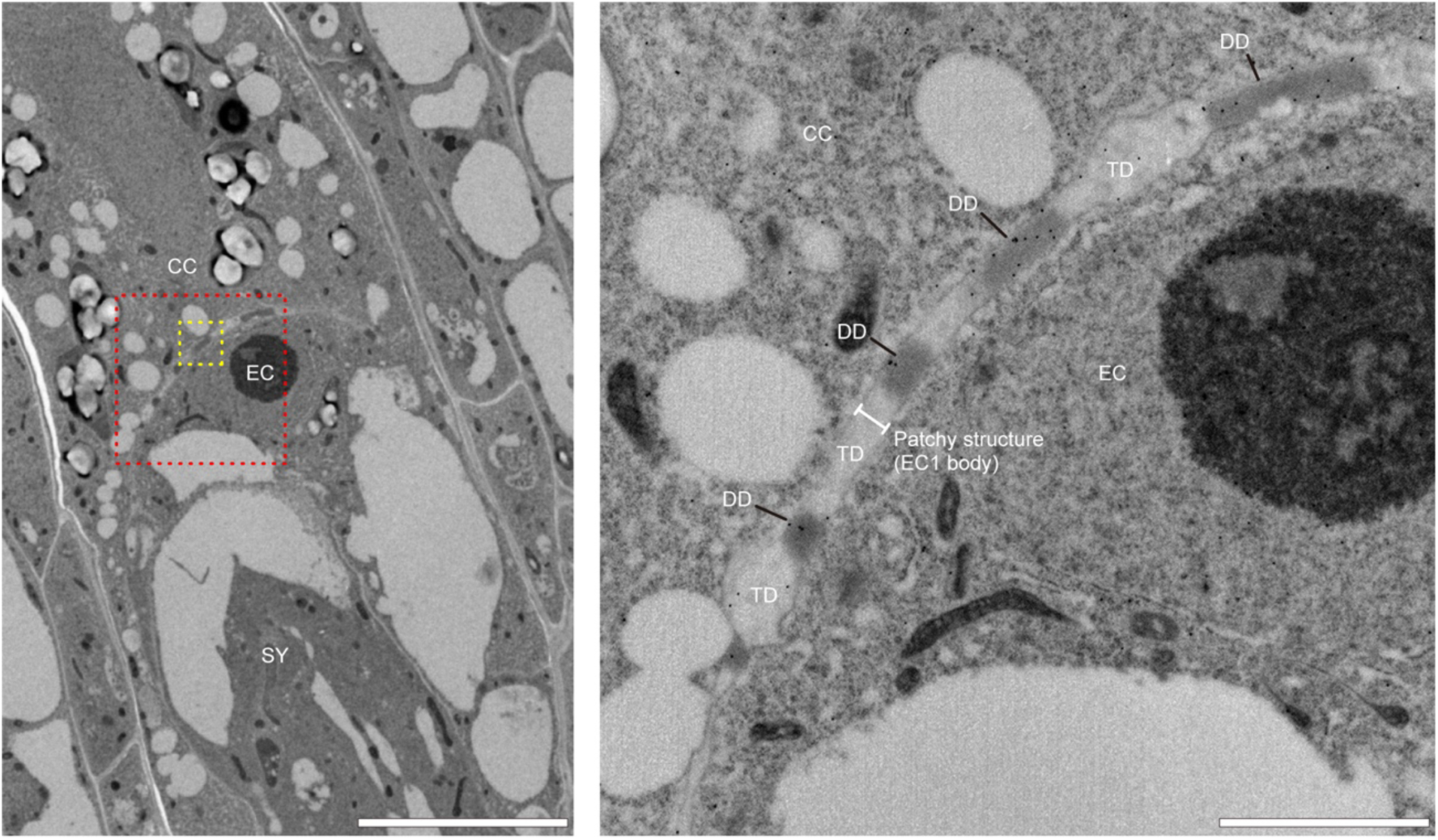
Immunogold staining of EC1.1 in mature ovules. Representative immunogold electron micrograph of a wild-type mature ovule using an anti-EC1.1 antibody. Magnified views of the red and yellow dashed boxes in the left panel are shown in the right panel and Fig. 3c, respectively. Abbreviations: EC, egg cell; CC, central cell; SY, synergid cell; DD, electron-dense domain; TD, translucent domain. Scale bars, 10 µm in the left panel; 2 µm in the right panel.

**Extended Data Fig. 5.**
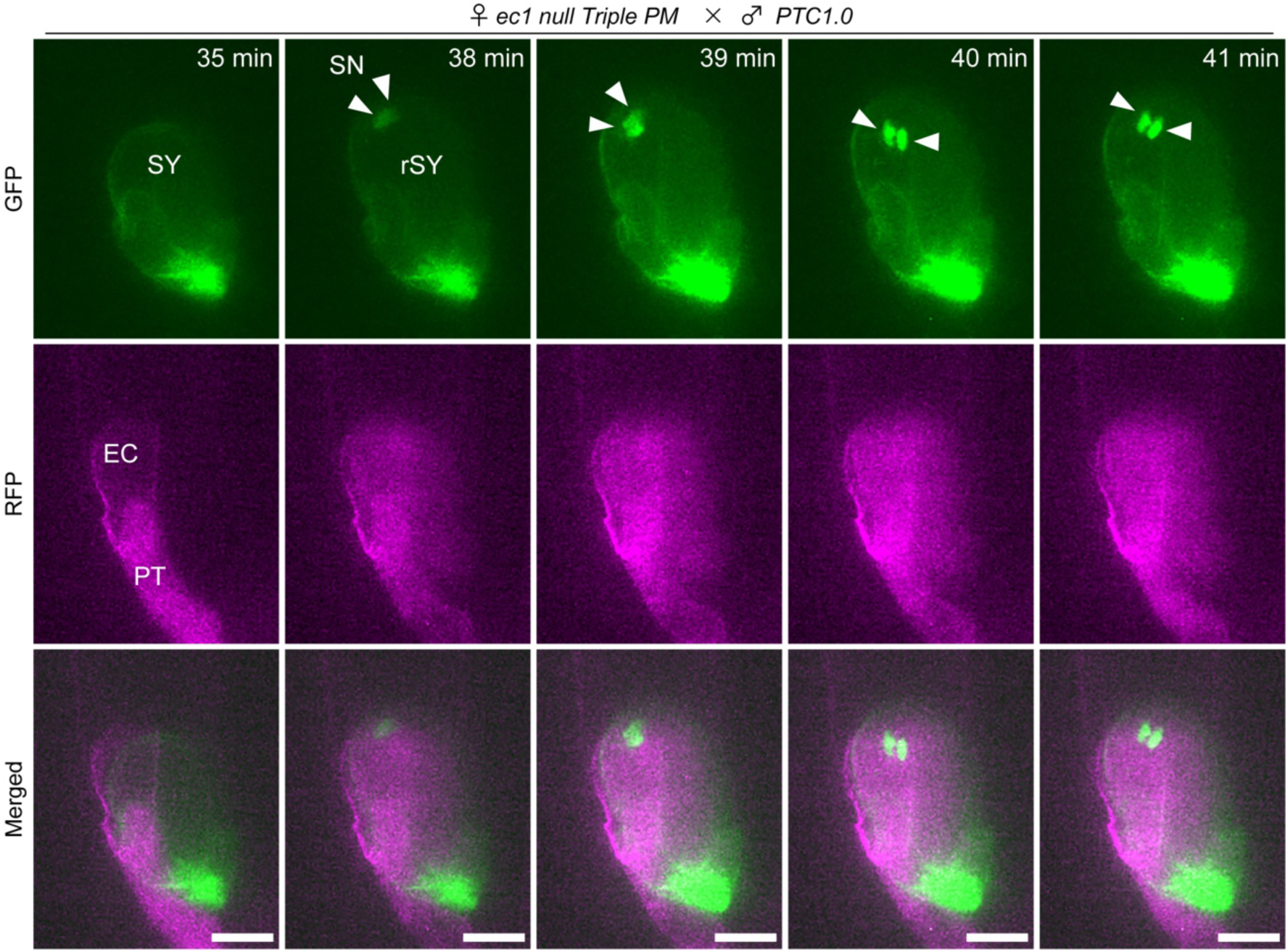
Pollen tube discharge in the *ec1 null* mutant. Semi-*in vivo* fertilization assay using ovules from the *ec1 null* mutant carrying the *Triple PM* marker and pollen from the *PTC1.0* marker line. Maximum-intensity projections of GFP, RFP, and merged channels acquired at five representative time points spanning pollen tube discharge are shown. Arrowheads indicate sperm nuclei (SN). See also Supplementary Video 8. Abbreviations: PT, pollen tube; EC, egg cell; CC, central cell; SY, synergid cell; rSY, receptive synergid. Scale bars, 10 µm.

**Extended Data Fig. 6.**
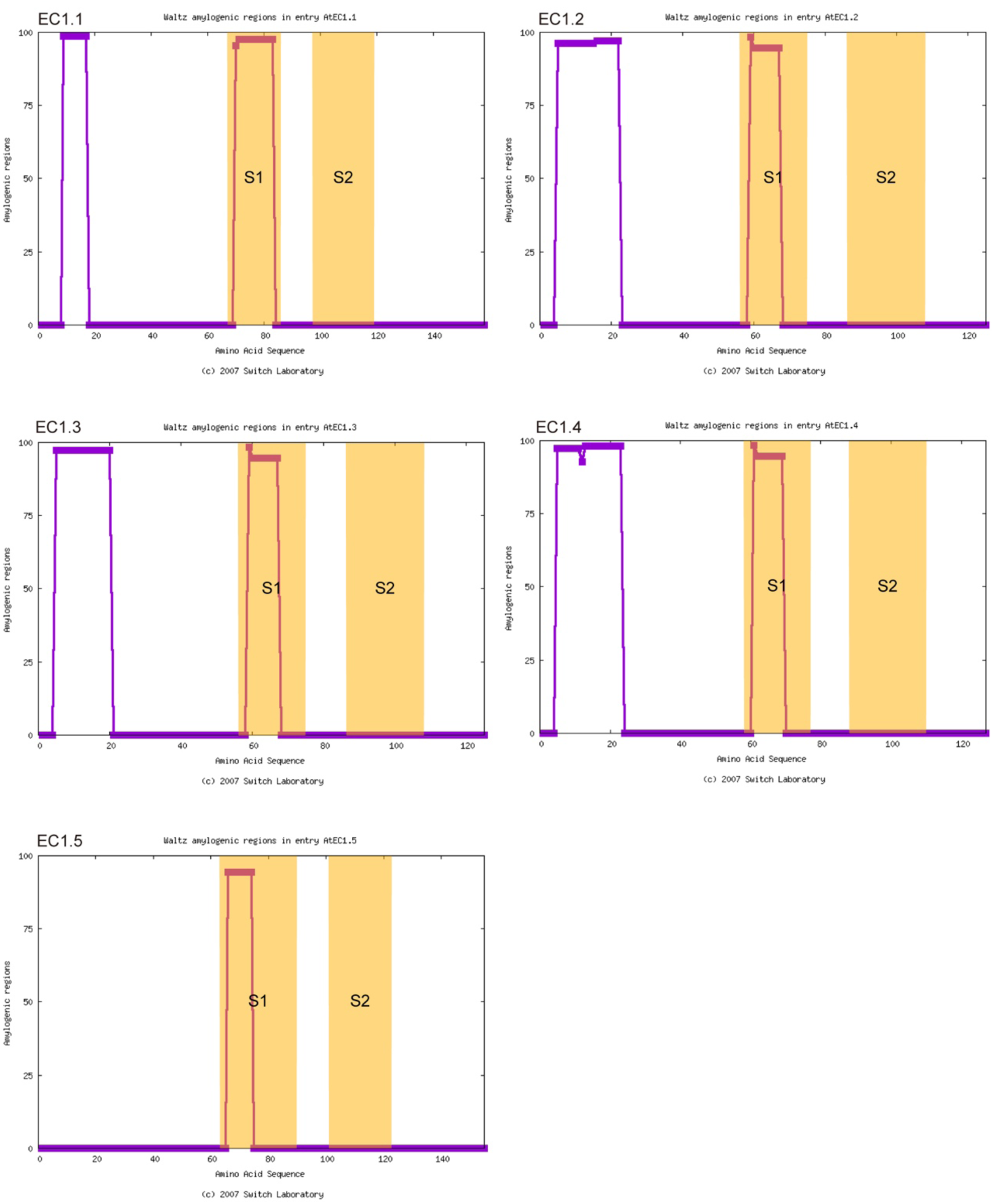
Prediction of amyloidogenic residues in EC1 peptides. Full-length amino acid sequences of *Arabidopsis* EC1.1–EC1.5 were analyzed using WALTZ, a predictor of amyloidogenic regions. Conserved S1 and S2 signature motifs are highlighted in yellow.

**Extended Data Fig. 7.**
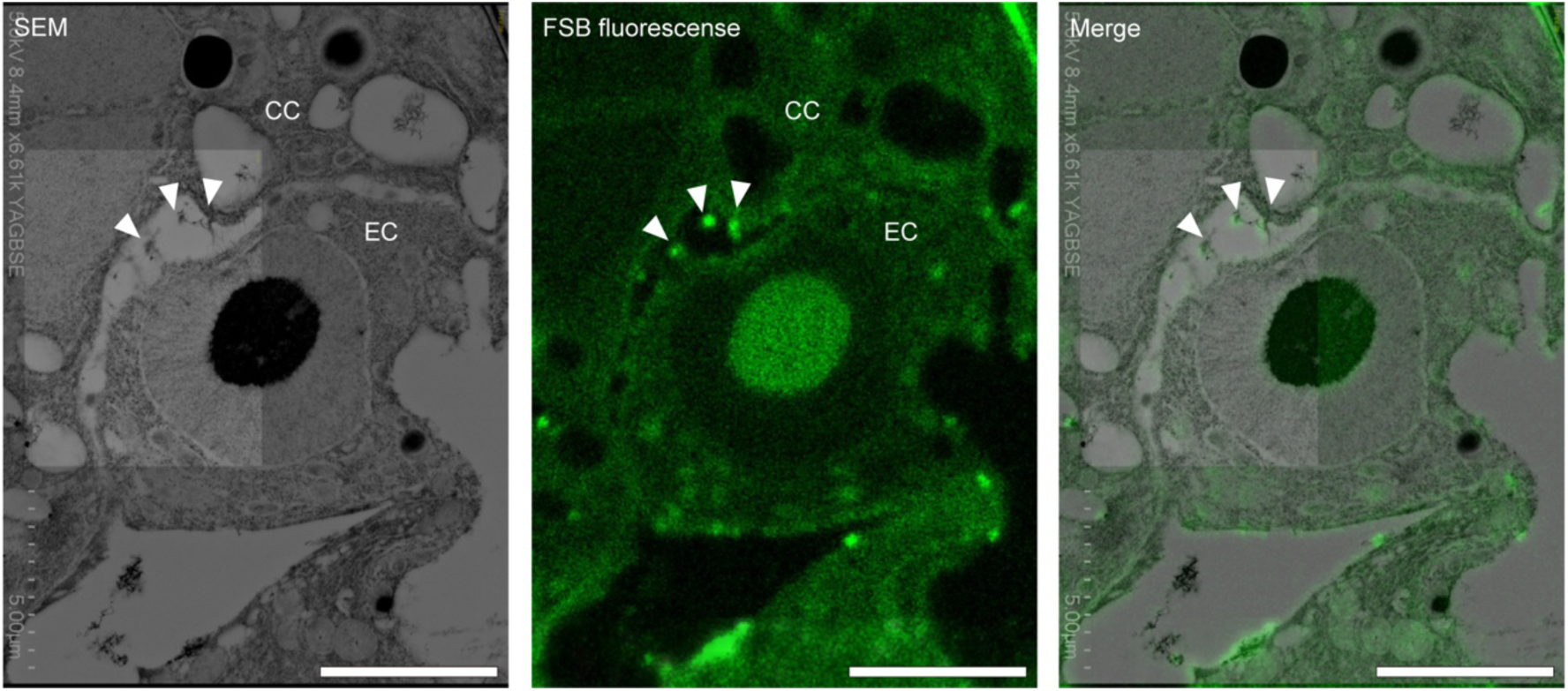
Colocalization of FSB-positive puncta with EC1 bodies. Correlative light and electron microscopy (CLEM) of mature ovule sections. FSB fluorescence was imaged before field-emission scanning electron microscopy (FE-SEM). Left, FE-SEM image. Middle, FSB fluorescence image. Right, merged image. Arrowheads indicate FSB-positive puncta corresponding to extracellular structures at the EC–CC interface. Abbreviations: EC, egg cell; CC, central cell. Scale bars, 5 µm.

**Extended Data Fig. 8.**
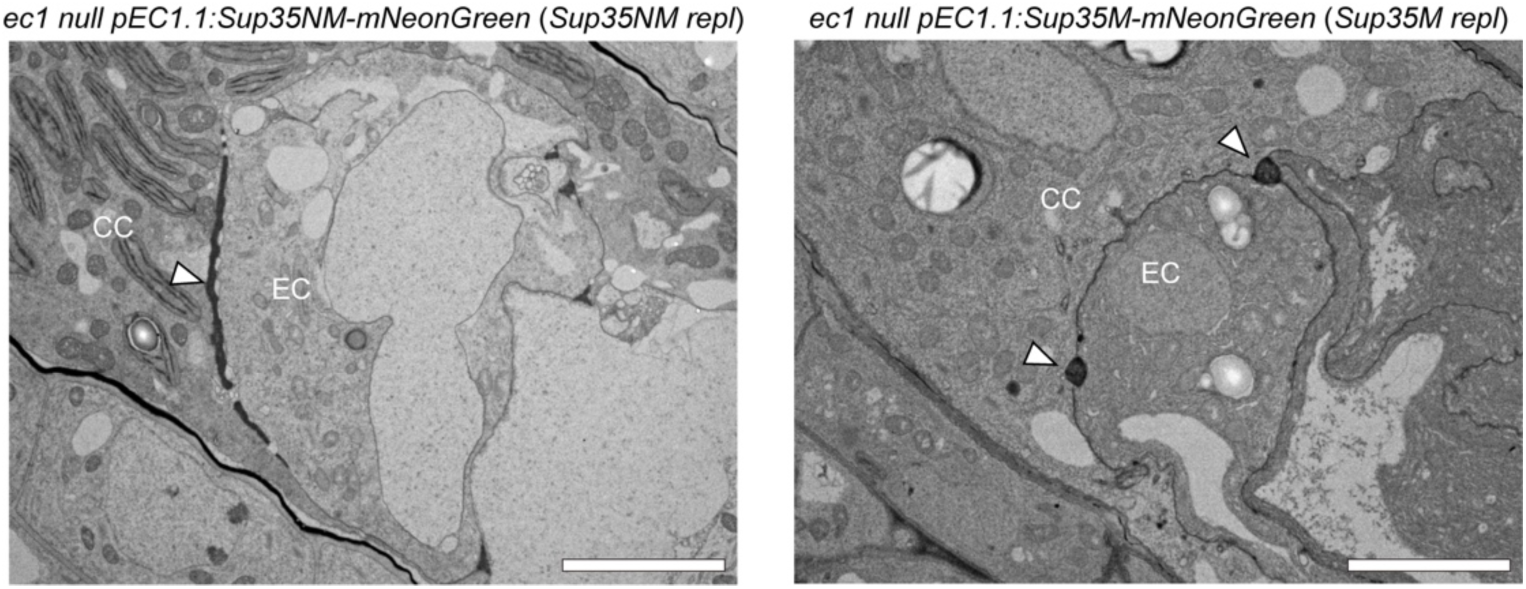
Electron micrographs of ovules expressing Sup35 replacement proteins. Representative transmission electron micrographs of ovules from *Sup35NM replacement* (*Sup35NM repl*) (n = 9) and *Sup35M replacement* (*Sup35M repl*) (n = 11) lines. Arrowheads indicate extracellular electron-dense deposits. Abbreviations: EC, egg cell; CC, central cell. Scale bars, 5 µm.

**Extended Data Fig. 9.**
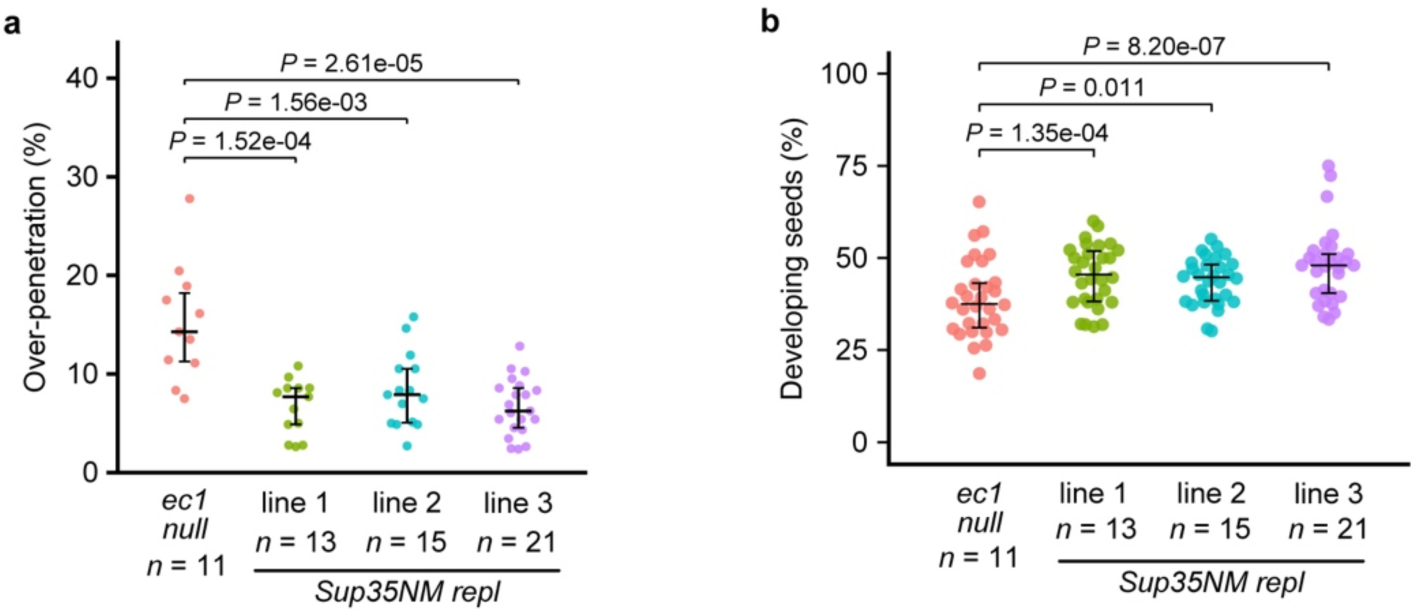
Phenotypic analyses of individual Sup35NM replacement lines. **a**, Frequencies of ovules exhibiting over-penetration in three independently generated *Sup35NM replacement* (*Sup35NM repl*) lines. **b,** Seed set of the corresponding lines. Combined data from the three *Sup35NM repl* lines shown in (**a**) and (**b**) are presented in Fig. 5d and Fig. 5f, respectively.

**Extended Data Fig. 10.**
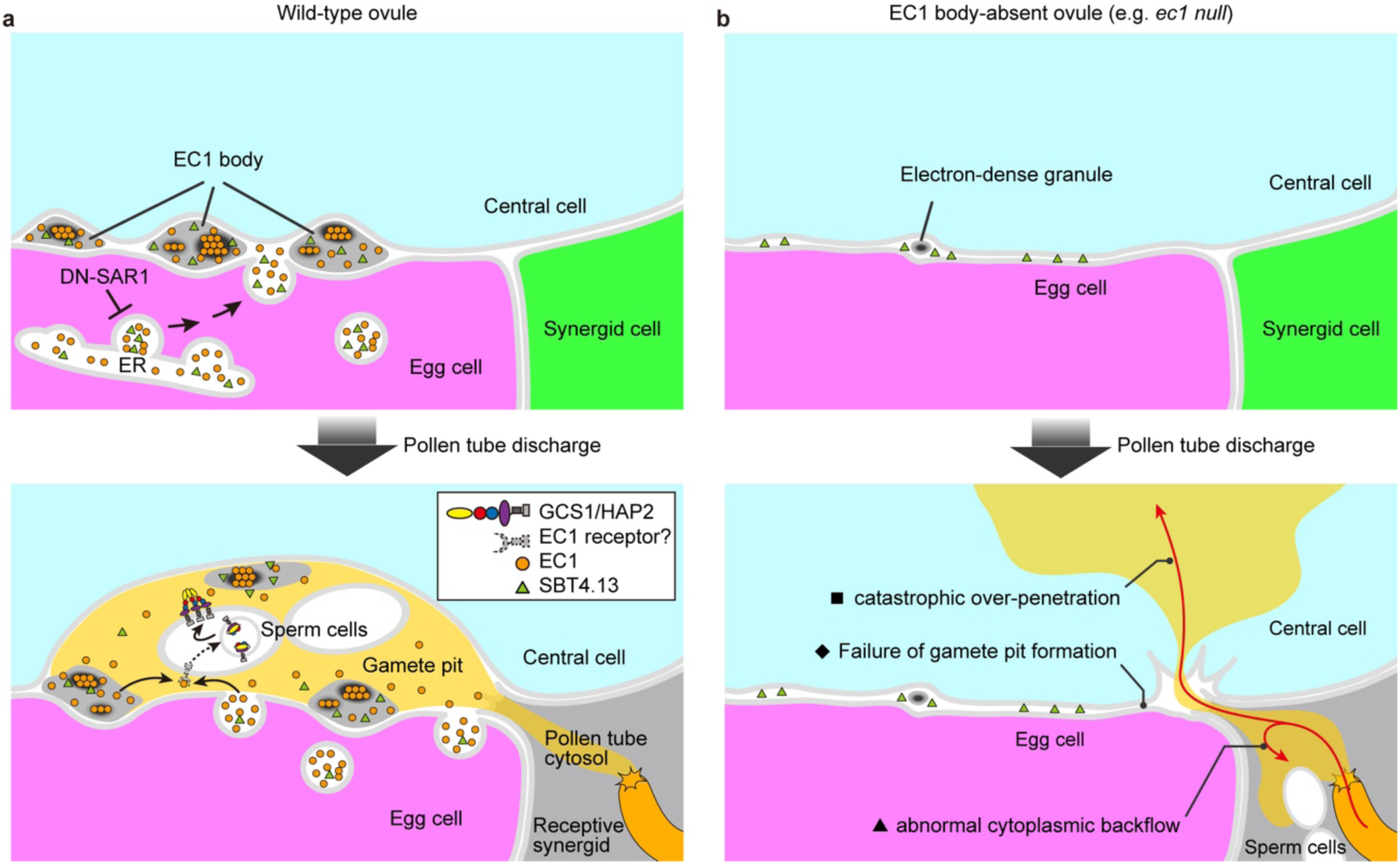
Model of EC1-mediated accurate sperm cell delivery. **a**, Schematic model of accurate sperm-cell delivery in wild-type ovules. EC1 peptides are secreted from the egg cell and assemble into EC1 bodies at the EC–CC interface through the secretory pathway, a process that requires ER-to-Golgi trafficking and is inhibited by dominant-negative SAR1 (DN-SAR1). EC1 bodies contain EC1 peptides together with other secreted proteins, including SBT4.13, and a subset of EC1 peptides forms amyloid-like assemblies (dark ellipses). Upon pollen tube discharge, rapid influx of pollen tube cytoplasm remodels the EC–CC interface in a cell-wall-independent manner, creating a transient gamete pit that enables accurate sperm-cell positioning. EC1 peptides supplied by diffusion from EC1 bodies together with continued secretion from the egg cell are proposed to activate GENERATIVE CELL SPECIFIC 1/HAPLESS 2 (GCS1/HAP2) relocation in released sperm cells through a putative EC1 receptor, thereby promoting successful double fertilization. **b,** Model of aberrant sperm-cell reception in ovules lacking EC1 bodies (for example, *pEC1.1:DN-SAR1* or *ec1 null*). In the absence of EC1 bodies, the EC–CC interface remains closely apposed except for a few residual electron-dense granules. Upon pollen tube discharge, pollen tube cytoplasm fails to remodel and open the EC–CC interface, thereby preventing gamete pit formation. As a consequence, released sperm cells are frequently displaced from the EC–CC interface by abnormal cytoplasmic backflow and remain trapped within the receptive synergid. In parallel, pollen tube contents can breach the central-cell plasma membrane, resulting in catastrophic over-penetration into the central-cell cytoplasm. Three characteristic defects are highlighted: ◆ failure of gamete pit formation, ▲ abnormal cytoplasmic backflow, ∎ catastrophic over-penetration. Together, these defects disrupt spatial confinement of pollen tube contents and accurate sperm-cell delivery, ultimately compromising double fertilization.

