## Supplementary Information for "EC1 body controls sperm cell reception *via* gamete pit formation"

**Captions for Supplementary Videos**

#### **Supplementary Video 1 | Dynamics of gamete pit formation.**

Semi-*in vivo* fertilization assay using ovules from the wild-type *Triple PM* marker line and pollen from the *PTC1.0* marker line. Time-lapse confocal imaging captures transient gamete pit formation immediately following pollen tube discharge. See also Fig. 1a,b. Scale bar, 10 µm.

**Supplementary Video 2 | Time-lapse imaging of pollen tube contents penetrating the receptive synergid.**

### Semi-*in vivo* fertilization assay using ovules expressing the *pMYB98:PIP2A-GFP* plasma membrane marker and pollen from the *PTC1.0* marker line. Time-lapse confocal imaging captures pollen tube contents penetrating the receptive synergid immediately following pollen tube discharge. See also Fig. 1c.

### **Supplementary Video 3 | Three-dimensional reconstruction of pollen tube contents penetrating the receptive synergid.**

### Three-dimensional reconstruction of a mature ovule expressing the *pMYB98:PIP2A-GFP* plasma membrane marker after reception of a *PTC1.0* pollen tube. The reconstruction was generated from Z-series confocal images acquired immediately after the time-lapse imaging shown in Fig. 1c and Supplementary Video 2.

**Supplementary Video 4 | Three-dimensional reconstruction of patchy extracellular structures.**

Three-dimensional reconstruction generated from serial FIB-SEM images of a mature unfertilized ovule. The egg cell (yellow), central cell (blue), synergid cells (light and dark green), and patchy extracellular structures (red) were manually segmented. The reconstruction reveals a network of patchy extracellular structures predominantly distributed along the EC–CC interface. See also Fig. 2b.

### **Supplementary Video 5 | Three-dimensional reconstruction of patchy extracellular structures in a live Arabidopsis ovule.**

### Three-dimensional reconstruction of a mature ovule expressing *pSBT4.13:SBT4.13-Clover*, revealing patchy extracellular structures at the EC–CC interface in living tissue. See also Extended Data Fig. 1b. Scale bar, 10 µm.

**Supplementary Video 6 | Abnormal pollen tube discharge in ovules with impaired COPII-mediated protein secretion in the egg cell.**

Semi-*in vivo* fertilization assay using ovules from the *pEC1.1:DN-SAR1 Triple PM* marker line and pollen from the *PTC1.0* marker line. Time-lapse confocal imaging captures defective pollen tube discharge accompanied by impaired sperm-cell positioning following pollen tube discharge. See also Extended Data Fig. 3. Scale bar, 10 µm.

**Supplementary Video 7 | Three-dimensional reconstruction of EC1.4-containing patchy extracellular structures in a live *Arabidopsis* ovule.**

Three-dimensional reconstruction of a mature ovule expressing *pEC1.4:EC1.4-Clover*, revealing patchy extracellular structures at the EC–CC interface in living tissue. See also Fig. 3a. Scale bar, 10 µm.

**Supplementary Video 8 | Abnormal cytoplasmic backflow following pollen tube discharge in the *ec1 null* mutant.**

Semi-*in vivo* fertilization assay using ovules from the *ec1 null Triple PM* marker line and pollen from the *PTC1.0* marker line. Time-lapse confocal imaging captures sperm cells initially approaching the EC–CC interface before being rapidly displaced toward the receptive synergid by abnormal cytoplasmic backflow following pollen tube discharge. See also Fig. 3e and Extended Data Fig. 5. Scale bar, 10 µm.

**Supplementary Video 9 | Apple-green birefringence of Congo Red-stained EC1.1 amyloid assemblies.**

Polarized-light microscopy of a C-DAG bacterial colony expressing the secreted mature EC1.1 protein. Rotation of the polarizer while maintaining the specimen and illumination conditions demonstrates the characteristic apple-green birefringence of Congo Red-positive amyloid assemblies. See also Fig. 4f. Scale bar, 10 µm.

**Supplementary Methods**

**Plasmid construction**

To produce pDM396, a destination vector carrying the *FWA* promoter, the 3,274-bp upstream from start codon of the *FWA* was amplified from Col-0 genomic DNA by a PCR using ‘FWA_HindIII_F’ and ‘FWA_HindIII_R’ as primers and introduced into HindIII site of pGWB501^1^. To generate the pDM349, a plasmid carrying the *pSBT4.13:SBT4.13-Clover*, we amplified a genomic sequence of the *SBT4.13* including 2,040-bp upstream of the start codon and protein coding region without stop codon by a PCR using ‘SBT_IF_F’ and ‘SBT_IF_R ’ as primers and Col-0 genomic DNA as a template, then introduced the sequence into SmaI site of the pPZP221-CloN^2^ by a homologous recombination using the In-Fusion HD Cloning Kit (Clontech). To produce pDM339, a plasmid harboring the *pEC1.1::EC1.1-Clover*, we performed PCR using Col-0 genomic DNA as a template and ‘EC1.1_SalI_F’ and ‘EC1.1_BamHI_R3’ as primers and introduced into SalI/BamHI site of the pPZP221-CloN. Similarly, we amplified *pEC1.2:EC1.2* to *pEC1.5:EC1.5* fragments using forward and reverse primers in Supplementary Table 1 with corresponding gene names and inserted them into SalI/BamHI site of the pPZP221-CloN to produce pDM340, pDM341, pDM342, and pDM343, respectively.

To generate the pDME108, an entry vector containing the *WT-SAR1*, SAR1 was amplified from Col-0 genomic DNA by a PCR using ‘SAR1_F’ and ‘SAR1_R’ as primers and introduced into the pENTR/D-TOPO (Invitrogen) by Gateway TOPO cloning. The pDME109, an entry vector containing the *DN-SAR1*, was constructed as follows. First, the *DN-SAR1* sequence was amplified as two DNA fragments with 26-bp overlap from Col-0 genomic DNA by independent PCRs using ‘SAR1_F’ and ‘Sar1_H74L_R’, or ‘Sar1_H74L_F’ and ‘SAR1_R’ as primer sets. Then, full-length *DN-SAR1* was amplified from the DNA fragments by a PCR using ‘SAR1_F’ and ‘SAR1_R’ as primers and introduced into the pENTR/D-TOPO (Thermo Fisher Scientific) by Gateway TOPO cloning. LR recombinations of the pDME108 or pDME109 with the MU2000 were performed using LR clonase II (Thermo Fisher Scientific) to produce pDM644 or pDM645, binary vectors containing the *pEC1.1:WT-SAR1* or *pEC1.1:DN-SAR1*, respectively. LR recombination of the pDME109 with the pSAN37^3^ was performed using LR clonase II to produce pDS075, a binary vector containing the *pDD65:DN-SAR1*. Similarly, LR recombination of pDME109 with the pDM396 were performed using LR clonase II to produce pDM407, a binary vector containing the *pFWA:DN-SAR1*.

To produce the *Triple PM* marker, we generated three entry clones carrying plasma membrane markers: the pOR082 (*Venus-SYP132*), the pOR080 (*mTFP1-SYP132*), and the pOR084 (*mRuby2-SYP132*). As described previously in the pOR084 construction^4^, the pOR080 and pOR082 were generated by the same procedure from two pENTR/D-TOPO vectors containing entire sequences of mTFP1^5^ or Venus^6^. LR recombinations between pOR082 with pDM286, pOR080 with pDM396, and pOR084 with MU2000 were performed to generate pDM429, pDM430, and pDM433, respectively. We then prepared three cassettes of plasma membrane markers for the *Triple PM* marker. The first cassette *pMYB98:Venus-SYP132* was amplified from pDM429 by a PCR using ‘pPZP221MYB98_F’ and ‘NOSter_Linker1_R’ as primers; the second cassette *pFWA:mTFP1-SYP132* was amplified from pDM430 by a PCR using ‘Linker1_FWA_F’ and ‘NOSter_Linker2_R’ as primers; and the third cassette *pEC1.1:mRuby2-SYP132* was amplified from pDM433 by a PCR using ‘Linker2_EC1.1_F’ and ‘pPZP221NOSter_R’ as primers. The three gene cassettes and SmaI-digested pPZP221 were assembled using the NEBuilder HiFi DNA Assembly Cloning Kit (New England Biolabs, MA, USA) to produce the Triple PM marker plasmid pDS270.

To construct pDS080, a pPZP221 binary vector carrying the *mNeonGreen* cDNA^7^, was constructed as follows. A 9,790-bp pPZP221 plasmid backbone was prepared by double-digesting pPZP221Ru^2^ with AscI and EcoRI to remove the mRUBY2 sequence. This backbone was then assembled with a 790-bp DNA fragment of the mNeonGreen cDNA, which was amplified by PCR using primers pPZP221_Asc1_mNG_F_366 and pPZP221_EcoR1_R_365. The two fragments were joined using the NEBuilder HiFi DNA Assembly kit. Plasmid pDS161, a binary vector containing the *EC1.1* promoter and signal sequence upstream of the *mClover* cDNA, was constructed via three-fragment assembly using NEBuilder. The assembly included: (1) an 8,985-bp DNA fragment obtained by HindIII digestion of pPZP221NosT; (2) a 2,228-bp fragment of the promoter and signal peptide coding region of the *EC1.1* amplified using primers pEC1.1_Hind3_NEB_F_493 and mClover_SP(EC1.1)_R_498; and (3) a 756-bp fragment of the mClover cDNA amplified using primers SP(EC1.1)_mClover_F_499 and pPZP221N_mClover_R_NEB_R_496. Plasmid pDS212, designed to express the *Sup35NM-mNG* under the control of the *EC1.1* promoter, was generated by assembling three fragments: (1) a 9,780-bp DNA fragment prepared by AscI digestion of pDS080; (2) a 2,236-bp fragment containing the *EC1.1* promoter and signal sequence using primers pEC1.1_Asc1_NEB_F_584 and SP(EC1.1)-Sup35NM_NEBR_585; and (3) an 820-bp fragment of the Sup35NM cDNA amplified from pVS72 using primers Sup35NM-SP(EC1.1)_NEB_F_586 and Sup35NM_Asc1_NEB_R_587. To generate *pEC1.1:Sup35M-mNG*, site-directed mutagenesis was performed using the QuickChange method^8^. PCR was conducted using pDS212 as a template with primers spEC1.1-SUP35M_F and spEC1.1-SUP35M_R. The original template DNA was then eliminated by DpnI digestion to yield the desired plasmid.　For amyloidogenicity assay, EC1.1–EC1.5 coding sequences lacking the signal peptide were cloned individually into the pVS105 export vector by insertion into the NotI and XbaI sites^9^. The resulting constructs were designated pDS191 (EC1.1), pDS192 (EC1.2), pDS193 (EC1.3), pDS194 (EC1.4) and pDS195 (EC1.5).

**Supplementary Table**

**Supplementary Table 1| Primers used in this study.**

| Primer name | Sequence (5' --> 3') |
| --- | --- |
| SBT_IF_F | CTCTAGAGGATCCCCAACATAACACATGTCACGAAC |
| SBT_IF_R | GCGCCCACCCTTCCCGTAATCACTAGTATAAACAACAATG |
| EC1.1_SalI_F | GCGGTCGACCCTGGAAAGCAAGAACAAAAGG |
| EC1.1_BamHI_R3 | CGCGGATCCCAGGGTTAGAAGGAGAAGCAG |
| EC1.2_SalI_F | GCGGTCGACGGGTTTAGAGAAAGACACACGG |
| EC1.2_BamHI_R3 | CGCGGATCCCAAGTTTCACAGAGGAAGGCGCC |
| EC1.3_SalI_F | GCGGTCGACCCAGACGGTTGCACTCCC |
| EC1.3_BamHI_R3 | CGCGGATCCCAAGTTTGACAGGGGAAAGAGC |
| EC1.4_SalI_F | GCGGTCGACGCTGATTAGAGCCGACGATGTC |
| EC1.4_BamHI_R3 | CGCGGATCCCAACTATTTTGGGAGACGGAGCC |
| EC1.5_SalI_F | GCGGTCGACTCATTTTTTCCAGACACGAATTG |
| EC1.5_BamHI_R3 | CGCGGATCCCATAATCAAGTCCGGGATACGTTATC |
| Sar1_H74L_F | GATTTGGGTGGCCtTCAGATTGCTCG |
| Sar1_H74L_R | CGAGCAATCTGAAGGCCACCCAAATC |
| Sar1_F | caccATGTTTTTATTCGATTGGTTCTATGG |
| Sar1_R | CTACTTGATATACTGAGATAGCC |
| FWA_HindIII_F | GCGAAGCTTGGTAGGCTAATAATCAGAAGCCCT |
| FWA_HindIII_R | GCGAAGCTTTCCCTCAATGCAATAACCTGGAC |
| pPZP221MYB98_F | CGACTCTAGAGGATCCCCGCGGCGGAGATAGTGGCTGAG |
| NOSter_Linker1_R | GGACAATGGTACCACATCCTCATTAGGCACCCCAGGCTTTAC |
| Linker1_FWA_F | GATGTGGTACCATTGTCCGGTAGGCTAATAATCAGAAGCCCT |
| NOSter_Linker2_R | CACCAACAACATTTTGTGCTCATTAGGCACCCCAGGCTTTAC |
| Linker2_EC1.1_F | CACAAAATGTTGTTGGTGTGCCTTATGATTTCTTCGGTTTC |
| pPZP221NOSter_R | CTACTCGAGATTGGTACCCACTAGCTCATTAGGCACCCCAGGCTTTAC |
| pPZP221_Asc1_mNG_F | CGGGAAGGGTGGGCGCGCCTCTGGAGGTGGAGGTTCAGGTGGAGGTGGAATGGTGAGCAAGGGCG |
| pPZP221_EcoR1_R | TAGTTTAATTAAGAATTCTCCACCTCCACCTGACTTGTACAGCTCGTCCATGC |
| pEC1.1_Hind3_NEB_F | AAAACGACGGCCAGTGCCAAGCTTCCTGGAAAGCAAGAACAAAAGG |
| mClover_SP(EC1.1)_R | CTCGCCCTTGCTCACCATGCGAGCTGTCACTGTGG |
| SP(EC1.1)_mClover_F | TCCACAGTGACAGCTCGCATGGTGAGCAAGGGCG |
| pPZP221N_mClover_R_NEB_R | GTCGACCTGCAGGCATGCAAGCTTTTACTTGTACAGCTCGTCCATG |
| pEC1.1_Asc1_NEB_F | GAGGATCCCCGGGAAGGGTGGGCCTGGAAAGCAAGAACAAAAGG |
| SP(EC1.1)-Sup35NM_NEB_R | TGTTGCCTTGGTTTGAATCCGAGCGAGCTGTCACTGTGGAG |
| Sup35NM-SP(EC1.1)_NEB_F | CTTCCTCCACAGTGACAGCTCGCTCGGATTCAAACCAAGGCAAC |
| Sup35NM_Asc1_  NEB_R | GAACCTCCACCTCCAGAGGCGCGGTGATGATGGTGATGGTGATCG |
| spEC1.1-SUP35M_F | CACAGTGACAGCTCGCTCTTTGAACGACTTTCAAAAGCAACAAAAG |
| spEC1.1-SUP35M_R | GCGAGCTGTCACTGTGGAGGAAGCCACCATGAGCATG |
